# An expanded toolkit for Drosophila gene tagging using synthesized homology donor constructs for CRISPR mediated homologous recombination

**DOI:** 10.1101/2021.12.24.474112

**Authors:** Oguz Kanca, Jonathan Zirin, Yanhui Hu, Burak Tepe, Debdeep Dutta, Wen-Wen Lin, Liwen Ma, Ming Ge, Zhongyuan Zuo, Lu-Ping Liu, Robert W. Levis, Norbert Perrimon, Hugo J. Bellen

## Abstract

Previously, we described a large collection of Drosophila strains that each carry an artificial exon containing a *T2AGAL4* cassette inserted in an intron of a target gene based on CRISPR-mediated homologous recombination (Lee *et al*., 2018). These alleles permit numerous applications and have proven to be very useful. Initially, the homologous recombination-based donor constructs had long homology arms (>500 bps) to promote precise integration of large constructs (>5kb). Recently, we showed that *in vivo* linearization of the donor constructs enables insertion of large artificial exons in introns using short homology arms (100-200 bps) (Kanca *et al*., 2019a). Shorter homology arms make it feasible to commercially synthesize homology donors and minimize the cloning steps for donor construct generation. Unfortunately, about 50% of Drosophila genes lack suitable coding introns for integration of artificial exons. Here, we report the development of new set of constructs that allow the replacement of the coding region of genes that lack suitable introns with a *KozakGAL4* cassette, generating a knock-out/knock-in allele that expresses GAL4 similarly as the targeted gene. We also developed custom vector backbones to further facilitate and improve transgenesis. Synthesis of homology donor constructs in custom plasmid backbones that contain the target gene sgRNA obviates the need to inject a separate sgRNA plasmid and significantly increases the transgenesis efficiency. These upgrades will enable the targeting of nearly every fly gene, regardless of exon-intron structure, with a 70-80% success rate.

## Introduction

The Drosophila Gene Disruption Project (GDP) aims to generate versatile genetic tools for most genes to facilitate the study of gene function *in vivo* and to create fly stocks for the community. The CRISPR mediated integration cassette (CRIMIC) approach is a recent addition to the GDP to target fly genes. The CRIMIC strategy is based on integrating a Swappable Integration Cassette (SIC) containing an artificial exon encoding *attP-FRT-Splice Acceptor (SA)-T2AGAL4-polyA-3XP3EGFP-polyA-FRT-attP* (T2AGAL4). The SIC is integrated in an intron between two coding exons (coding intron) by CRISPR mediated homologous recombination (Lee *et al*., 2018; Gnerer *et al*., 2015; Diao *et al*., 2015). The viral T2A sequence leads to the truncation of the nascent target gene polypeptide and re-initiation of translation of the downstream GAL4 as an independent protein. This cassette typically creates a strong loss of function allele of the targeted gene and expresses the yeast GAL4 transcription factor in a similar spatial and temporal pattern as the protein encoded by the targeted gene (Lee et al., 2018). These alleles can be used to: 1) determine the gene expression pattern; 2) study the effect of loss of function of the gene product; 3) replace the SIC through Recombinase Mediated Cassette Exchange (RMCE) (Bateman *et al*., 2006; Venken *et al*., 2011) with an artificial coding exon that encodes a fluorescent protein to assess protein subcellular localization (Venken et al., 2011) and identify interacting proteins ; 4) express UAS-cDNAs of the targeted gene and its variants to assess rescue of the mutant phenotype and conduct structure/function studies (Wangler *et al*., 2017); 5) excise the insert with UAS-Flippase to revert the phenotype (Lee *et al*., 2018).

The introduction of an artificial exon is only feasible for genes that contain a suitable large coding intron, typically 100 nt or more. This requirement makes nearly half of the genes inaccessible to strategies based on the use of artificial exons (Supplementary table 1; Figure 1A). In addition, the genes that do not have a suitable intron are typically smaller in size than genes that contain a suitable intron, and usually have fewer previously isolated, publicly available alleles than larger genes with a suitable intron.

**Figure 1.**
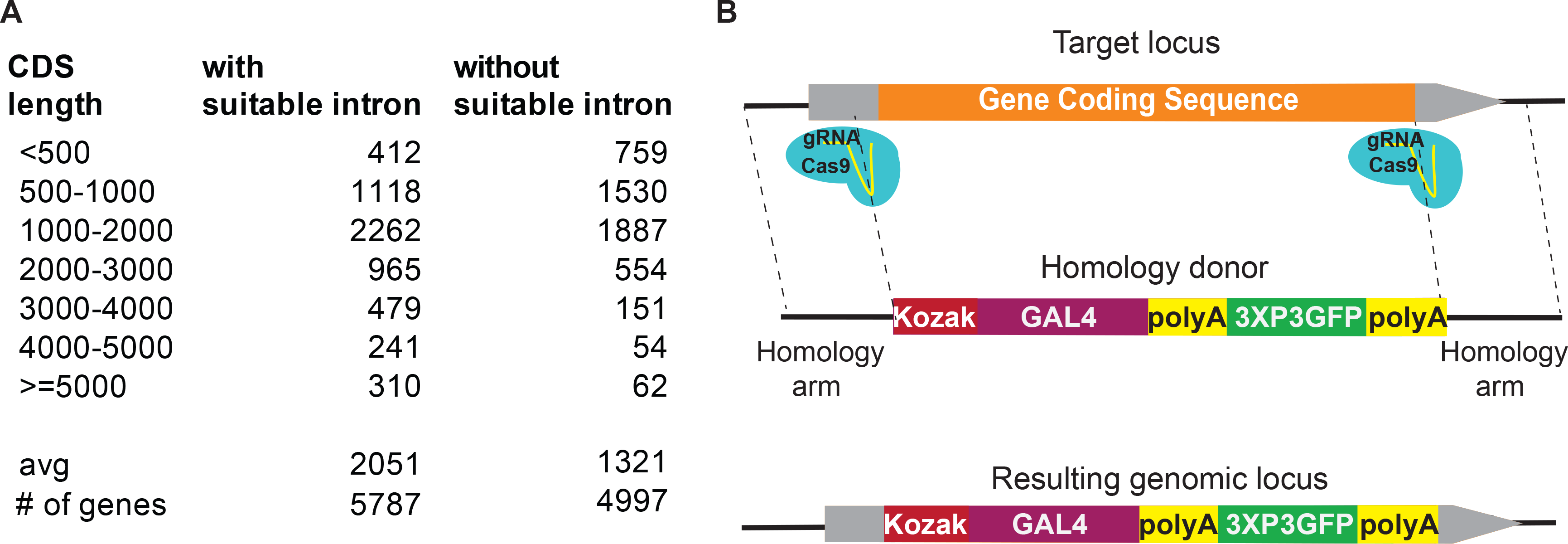
KozakGAL4 strategy can be used to generate GAL4 gene trap alleles for approximately 50% of Drosophila genes. (A) Analysis of Drosophila genome indicates that about half of the Drosophila genes do not have a suitably large coding intron for insertion of a T2AGAL4 cassette. These genes are on average shorter and have fewer genetic reagents compared to the genes that have a suitably large coding intron for inserting T2AGAL4 cassette. (B) Schematics of the KozakGAL4 targeting. Gray boxes, UTRs; orange box, gene coding region.

Here, we describe the development of a knock-out/knock-in strategy to replace the coding sequence of genes with a *Kozak sequence-GAL4 polyA-FRT-3XP3GFP- polyA-FRT* (*KozakGAL4*) cassette to target genes that lack introns that are suitable for artificial exon knock-ins. The targeted gene is cut by Cas9 using two sgRNAs, one targeting the 5’ untranslated region (UTR) and the other the 3’ UTR. We targeted ∼100 genes with this strategy and show that about 80% of the integrated KozakGAL4 cassettes lead to *UAS-mCherry* expression in the 3^rd^ instar larval brain, a ratio that is similar to what was observed for the T2AGAL4 strategy (Lee *et al*., 2018).

We also improved the design of the homology donor constructs that can be used for integration of either *KozakGAL4* or *T2AGAL4* cassettes. The use of short homology arms allows commercial DNA synthesis of the entire gene-specific portion of the donor plasmid, a cheaper and more efficient option than PCR-amplification and cloning of each homology arm (Kanca *et al*., 2019a). To further extend this approach, we developed a method in which the DNA sequences directing the transcription of the target gene-specific sgRNA(s) are synthesized together in the same segment as the homology arms. The design allows gene specific sgRNA to be synthesized together with the homology arms, eliminating the need to clone and inject a separate vector for the sgRNAs. We tested our new designs on ∼200 genes and show that the upgrades result in a transgenesis efficacy of ∼80%. The strategies that we introduce here allow targeting of nearly every gene in the fly genome, further streamline the generation of homology donor DNAs, increase efficiency as compared to previous strategies, and improve the rate of precise genome editing.

## Results and Discussion

### The KozakGAL4 cassette

Integration of an artificial exon is only feasible for genes that have coding introns large enough to identify an sgRNA target site that is located at a sufficient distance from the preceding splice donor and the following splice acceptor site. Based on our experience, an intron should be larger than 100 nt to be suitable for integration of an artificial exon. An analysis of the Drosophila genome shows that 5787 out of 13,931 protein-coding genes have a sufficiently large coding intron that is shared among all the annotated splicing isoforms of the gene. However, 8144 genes lack such introns, making them inaccessible for the T2AGAL4 and other artificial exon-based strategies. Genes with a suitable intron typically have larger coding sequences (2051 nt vs 1321 nt) and have a larger number of previously isolated mutant alleles based on FlyBase data (10.4 vs 3.4) than genes without a suitable intron(s) (Figure 1A, Supplementary table 1; Larkin *et al*., 2020). To integrate a GAL4 cassette that can be used in a similar manner as the *T2AGAL4* insertions, we developed the *KozakGAL4* knock-in/knock-out strategy. Kozak sequence is an optimal translation initiation site in eukaryotic mRNAs, and it is identified as (C/A) AA (C/A) AUG in Drosophila (Cavener, 1987; Kozak, 1986). We use CAAA as a Kozak sequence upstream of the start codon of GAL4. To replace the coding region of genes, we typically identify sgRNA target sites in the 5’ UTR and 3’ UTR (Figure 1B). To retain possible gene expression regulation by the 5’UTR, we select the upstream sgRNA target site that is closest to the start codon and that is not predicted to have off-target activity based on CRISPR Optimal Target Finder (Gratz *et al*., 2014). The location of the downstream sgRNA target site in the 3’UTR is less stringent since the endogenous 3’UTR is not included in the final transcript due to the *polyA* signal in the *KozakGAL4* cassette. The median 5’ and 3’ UTR lengths for *Drosophila* genes are 214 and 224 bps respectively which are typically large enough to identify putative sgRNA targets (Chen *et al*., 2011; Jan *et al*., 2011). In our experience the 5’UTR typically contains multiple sgRNA targets whereas 3’UTRs contain few candidate sgRNA target sites due to their A/T rich nature. In cases where a suitable sgRNA target site cannot be found in the 3’UTR, we target a site within the coding region, close to the stop codon, to minimize the coding region of the gene that remains. In cases where a suitable sgRNA site cannot be found within the 5’UTR region, the search is expanded to the promoter region and the sequence between the gRNA cut site and transcription start site is added to the homology region. In such cases, a single nucleotide substitution to eliminate the PAM sequence is introduced in the homology donor construct, preventing cutting of the homology donor.

We also developed alternative strategies to target genes without suitable introns and for which no proper sgRNA site could be identified within the 5’UTR or promoter. We generated the *SA-KozakGAL4-polyA-3XP3EGFP-polyA* cassette that can be introduced in an intron within the 5’UTR (Figure 1 Supplementary Figure1A).

Alternatively, for genes with small coding introns, we make two cuts: one within a coding intron just upstream of the SA of an exon; the second in the 3’UTR. The excised sequence is then replaced with a T2AGAL4 cassette (Figure 1 Supplementary Figure 1B).

### KozakGAL4 alleles drive expression of UAS-transgenes

There are two main approaches to generate alleles that express GAL4 that are not based on the T2AGAL4-based strategies. The first one is based on an insertion of minimal promoter-GAL4 coding sequences in a transposon backbone. The strategy is called enhancer trapping and was based originally on *GFP* and *LacZ* rather than *GAL4* (O’Kane and Gehring, 1987; Bellen *et al*., 1989). Upon mobilization of the transposon, lines are established where the GAL4 expression pattern is of interest (Brand and Perrimon, 1993; Lukacsovich *et al*., 2001; Hacker *et al*., 2003; Gohl *et al*., 2011). Given that they are inserted in the genome by transposable elements they are not always optimally placed to report the full expression pattern of the gene (Spradling *et al*., 1995; Mayer *et al*., 2013) but they have been used extensively as many reflect the expression pattern of a nearby gene (Wilson et al., 1989). Many however are not mutagenic (Spradling *et al*., 1999). The second strategy to generate alleles that may express GAL4 in the expression pattern of a gene is to clone a 500bps-5kb region upstream of the promoter of the gene upstream of the GAL4 coding sequences and inserting the transgene in the genome. There are large collections of these enhancer-GAL4 alleles and most aim to report the expression of enhancer fragments rather than reporting the expression pattern of the gene from which they are derived (Jenett *et al*., 2012; Manning *et al*., 2012; Pfeiffer *et al*., 2008). Hence, the available approaches can now be complemented with the KozakGAL4 approach that should incorporate all or most upstream regulatory information and generate a null allele of the targeted gene. The latter greatly facilitates rescue experiments using UAS-cDNA transgenes.

To assess the KozakGAL4 strategy, we targeted 109 genes to date and successfully replaced the coding region of 82 genes with the *KozakGAL4* cassette (Supplemental Table 2). We crossed 57 of these with *UAS-CD8mCherry* transgenic flies to determine the GAL4 expression of the targeted gene in the brain of wandering third instar larvae. Our previous findings, using *T2AGAL4* alleles have shown that ∼80% of all *T2AGAL4* alleles lead to specific expression in third instar larval brains (Lee *et al*., 2018). Similarly, with *KozakGAL4* alleles, we detected specific GAL4 expression for about 80% (46/57) of the genes (Figure 2A). Although *KozakGAL4* targeted genes are typically small, limiting the possible regulatory information in the coding region, it is possible that excision of coding and some UTR sequences may remove part of the regulatory input. We therefore tested whether a few *KozakGAL4* alleles drive the expression of the *UAS-CD8mCherry* in a similar pattern as the targeted gene. We selected a *KozakGAL4* allele that drives expression of the reporter in a restricted group of cells in the 3^rd^ instar larval brain and analyzed the single cell RNA sequencing (scRNAseq) data for the 3^rd^ instar larval brain to determine cell clusters that express the targeted gene (Ravenscroft *et al.,* 2020). We then used the same scRNAseq dataset to determine other genes expressed in overlapping clusters and that we previously targeted with *T2AGAL4*. *KozakGAL4* driven *UAS-CD8mCherry* reporter is expressed in a very similar expression pattern compared to *T2AGAL4* driven reporter expression of the genes that we identified through scRNAseq. The other genes expressed in the overlapping cluster according to scRNAseq, such as *serp*, *verm* and *emp* suggest that this cluster corresponds to tracheal cells, which is in line with the observed expression pattern through imaging (Figure 2B, Figure 2 Supplementary Figure 1B; Luschnig *et al.,* 2006; Lee *et al*., 2018). Comparison of the expression patterns of the other tested *KozakGAL4* targeted genes and *T2AGAL4* targeted genes that are expressed in overlapping cell groups showed overlapping expression patterns based on imaging as well (Figure 2 Supplementary Figure 1A). Hence, the use of scRNAseq data can provide an independent means of verification of accuracy of the observed reporter expression patterns.

**Figure 2.**
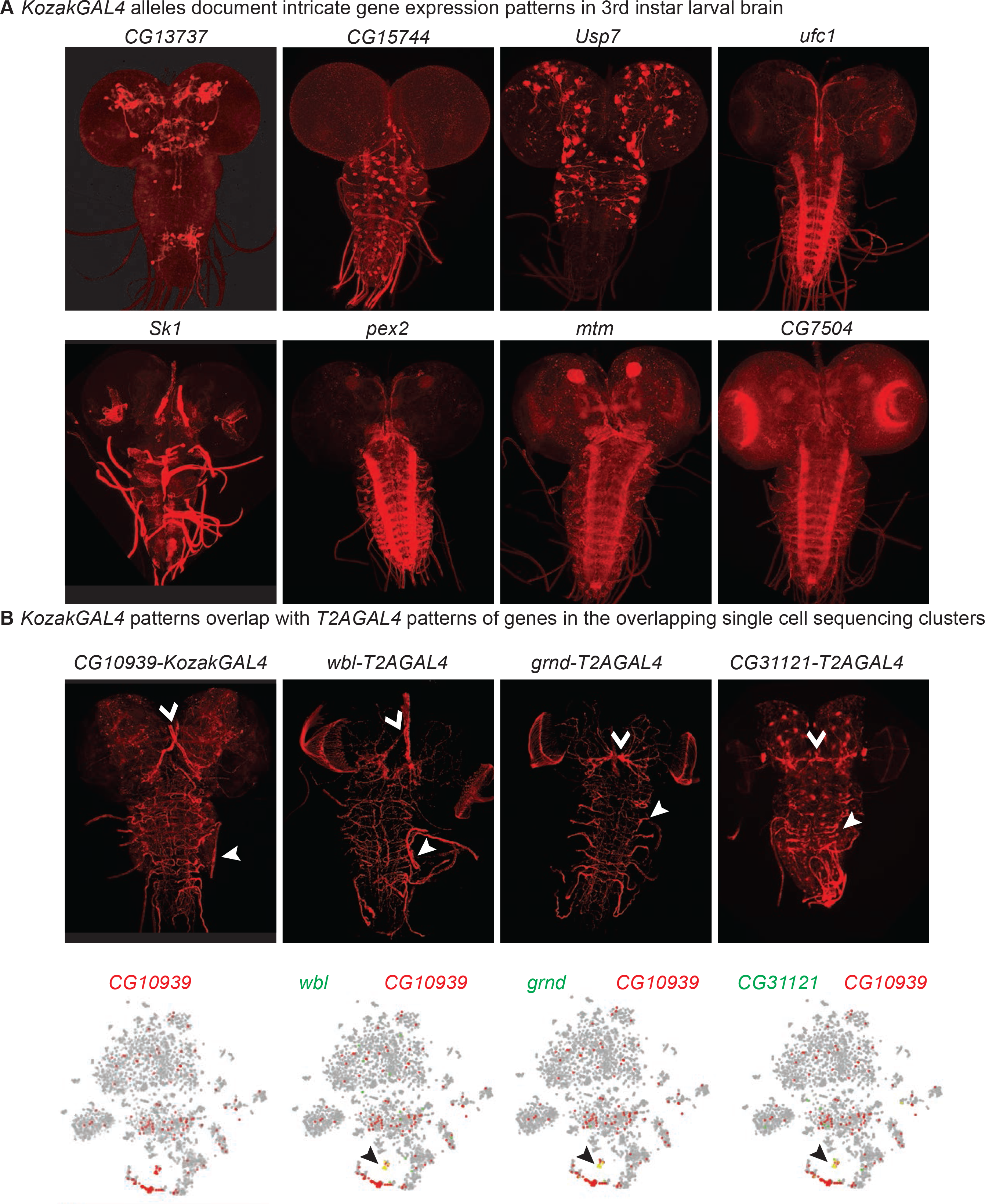
KozakGAL4 alleles document intricate gene expression patterns in third instar larval brains. (A)Examples of third instar larval brain gene expression patterns obtained by crossing KozakGAL4 allele of indicated genes with *UAS- CD8mCherry* flies. (B) The imaging results of reporter expression generated with *KozakGAL4* allele were compared to the expression pattern of genes that are expressed in similar cells by analysis of single cell sequencing data imaged using *T2AGAL4* alleles. Images are taken by crossing the *GAL4* alleles with *UAS- CD8mCherry.* Arrowheads point to the shared expression pattern.

Next, we determined if UAS-human or fly cDNAs could rescue the phenotype associated with 11 gene deletions caused by the *KozakGAL4* knock-ins. For *pngl*, *Wdr37, Tom70, CG8320, CG16787* and *IntS11* a UAS-fly or human cDNA rescued the *KozakGAL4* induced phenotypes, suggesting that the *KozakGAL4* is expressed where the targeted gene product is required for the gene function. For *pex2, pex16, fitm, PIG-A* and *CG34293* the expression of orthologous human cDNA did not rescue the associated phenotypes.

In summary, *KozakGAL4* offers a new means to disrupt gene function while expressing the GAL4 in the expression domain of the targeted genes. This approach allows us to tag the remainder of the genes that do not contain a suitable coding intron for the T2AGAL4 strategy which corresponds to 58% of all the genes.

### New vector backbones for synthesis of homology donor constructs that are also templates for sgRNA expression

We previously showed that linearizing the homology donor constructs *in vivo* allows for integration of large constructs in the genome through CRISPR-mediated homologous recombination even using short homology arms (Kanca *et al.,* 2019a). This approach makes inexpensive commercial synthesis of homology donor intermediates feasible. The intermediate vectors can be used for a single step directional cloning of the SIC in the homology donor intermediate vector. This greatly facilitates the generation of homology donor vectors which previously required four-way ligations with large homology arms. Moreover, this eliminates cloning failures (∼20-30%) and troubleshooting associated constructs with large homology arms (Kanca *et al*., 2019a). The resulting new homology donor vectors were previously injected together with two vectors that express two sgRNAs (pCFD3, Port *et al*., 2014) in embryos that express Cas9 in their germline. The first sgRNA targets the homology donor vector backbone to linearize the homology donor and does not have a target in the Drosophila genome (sgRNA1, Garcia-Marques *et al*., 2019). The second sgRNA vector expresses the sgRNA to target the gene and introduce the double strand DNA break that serves as a substrate for homologous recombination. In Kanca *et al*. (2019a) we demonstrated that injection of these constructs resulted in transgenesis efficiencies of about 60%.

We developed new approaches to increase the transgenesis efficiency of the custom DNA backbones, decrease the workload, and to simplify the generation of homology donor constructs. The first custom vector backbone that we tested has the U6- 3::sgRNA1 sequence in the vector backbone and sgRNA1 targets, on either side of the EcoRV site where the synthesized fragments are directionally integrated (vector backbone named pUC57_Kan_gw_OK, design named int200, Figure 3A). With this design, the homology donor vector intermediates that are commercially synthesized contain the sgRNA1 coding sequence, obviating the need to co-inject one of the sgRNA vectors. Having the sgRNA1 coding region in the backbone also helps with *in vivo* linearization of the homology donor since the homology donor construct and the sgRNA1 are delivered together in a single vector. The int200 design also removes the sgRNA1 target sites from the synthesized region as they are present in the vector. This allows increasing the homology arm length to 200 bps without increasing the cost of synthesis.

**Figure 3.**
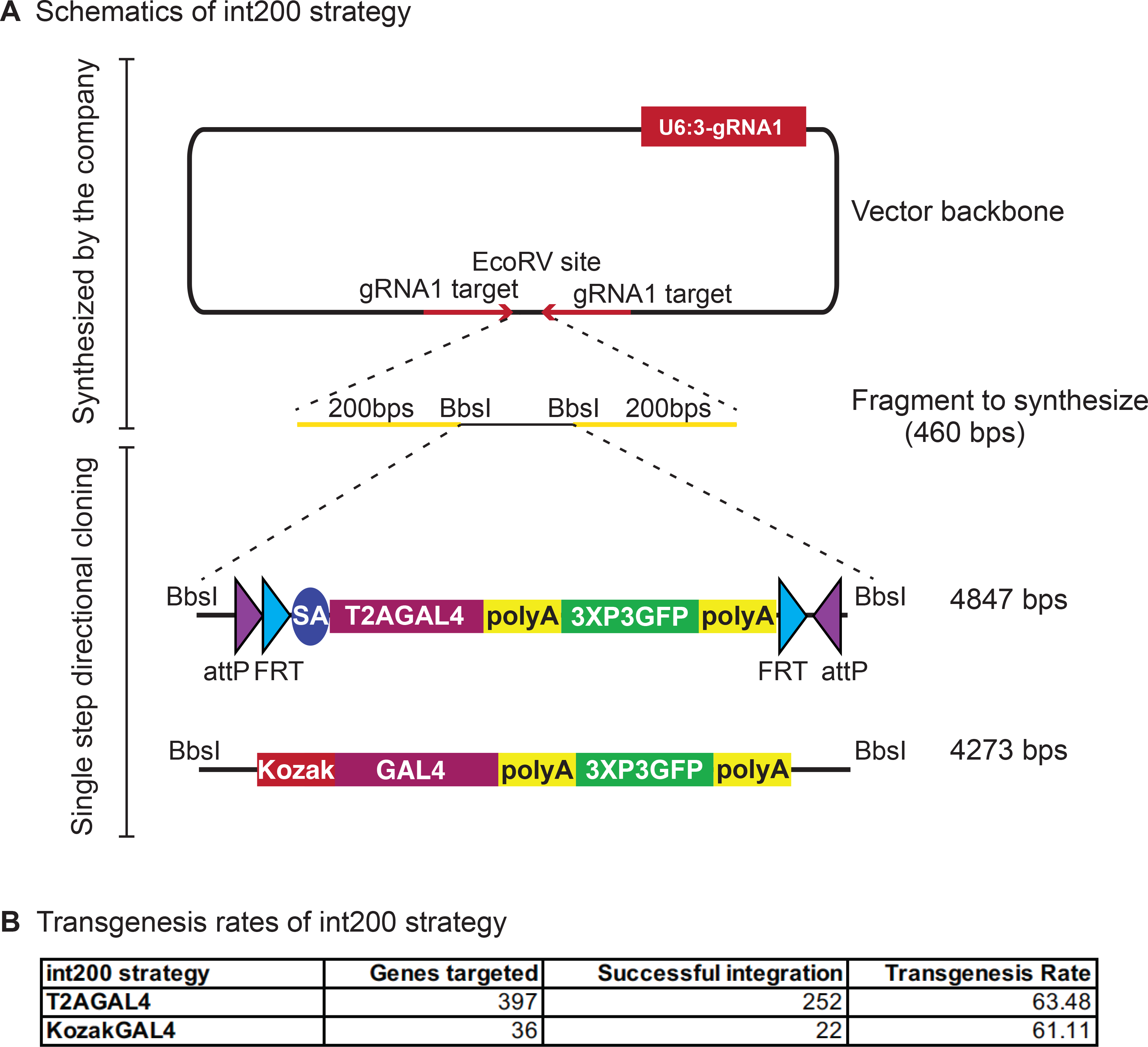
int200 strategy results in similar transgenesis success rates as the long homology arms CRIMICs. (A) Schematics of the int200 strategy. (B) Transgenesis data using int200_T2AGAL4 or int200_KozakGAL4 strategies.

We tested this int200 design for 397 genes with T2AGAL4 cassettes and 36 genes with KozakGAL4 cassette. For each construct, we injected 400-600 embryos that express Cas9 in the germline. For inserting the KozakGAL4 cassette, the two gene specific sgRNAs were cloned into pCFD5 (Port and Bullock, 2016). For both T2AGAL4 and KozakGAL4 insertions the int200 homology donor plasmid was co-injected with the plasmid that encodes the target specific sgRNA (pCFD3 for the former and pCFD5 for the latter case). We successfully integrated T2AGAL4 cassette in 252 genes (∼65% success rate) and replaced the coding region of 22 genes with KozakGAL4 (∼61% success rate) (Figure 3B). PCR verification of the inserts was performed by using gene specific PCR primers outside the homology region pointing towards the insert and a construct specific PCR primer. For 88% of the T2AGAL4 inserts we obtained PCR verification on both sides of the insert and for the remaining 12% we obtained PCR products on one side of the construct. For 91% of KozakGAL4 inserts we obtained amplicons on both sides of the insert. For the inserts with a single PCR verification, we sequenced the amplicon to ensure the insert is in the proper locus. Hence, the overall transgenesis success rate of the int200 method is about 65% (Figure 3B). This is very similar to the injection success rate of homologous recombination using large (0.5-1kb) homology arms (1165 insertions in 1784 targeted genes, Lee *et al*., 2018) but leads to very significant reductions in labor and cost. Additionally, int200 facilitates the cloning of homology donor constructs and eliminates cloning failures which reduce the overall successful targeting rate using large homology arms to ∼50% (successful cloning of 80% constructs that are injected with 65% transgenesis success rate). In summary, the int200 method provides a ∼30% gain in overall efficiency (from 50% to 65%).

To further optimize the custom vector backbone we repositioned the U6- 3::sgRNA1and added a partial tRNA construct directly upstream of the EcoRV site that is used to insert the synthesized fragments (Figure 4A). The partial tRNA allows adding the gene-specific sgRNA sequence (vector named pUC57_Kan_gw_OK2 and design named gRNA_int200 for T2AGAL4 constructs and named 2XgRNA_int200 for KozakGAL4 constructs). Hence, two or three sgRNA can be produced from the single injected plasmid. One of the sgRNA1 target sites is added to the synthesized fragment before the start of the homology arm and the other sgRNA1 target site is added to the backbone just downstream of the EcoRV site where the synthesized fragment is directionally inserted. This design obviates the need to clone a separate sgRNA vector to target the genomic locus. It also ensures simultaneous delivery of all the components of the homologous recombination reaction as they are delivered on a single plasmid. We have targeted 127 genes with gRNA_int200_T2AGAL4 donor plasmids (Supplemental Table 2) and successfully inserted the T2AGAL4 cassette in 95 genes (∼75% success rate, Figure 4B). We also tested whether genes for which the tagging failed using the int200 strategy (Figure 3) could be targeted with the gRNA_int200_T2AGAL4 using the same gene-specific sgRNA and homology arms. For 3 out of 4 genes tested, use of the gRNA_int200 strategy resulted in successful integration of T2AGAL4 cassette.

**Figure 4.**
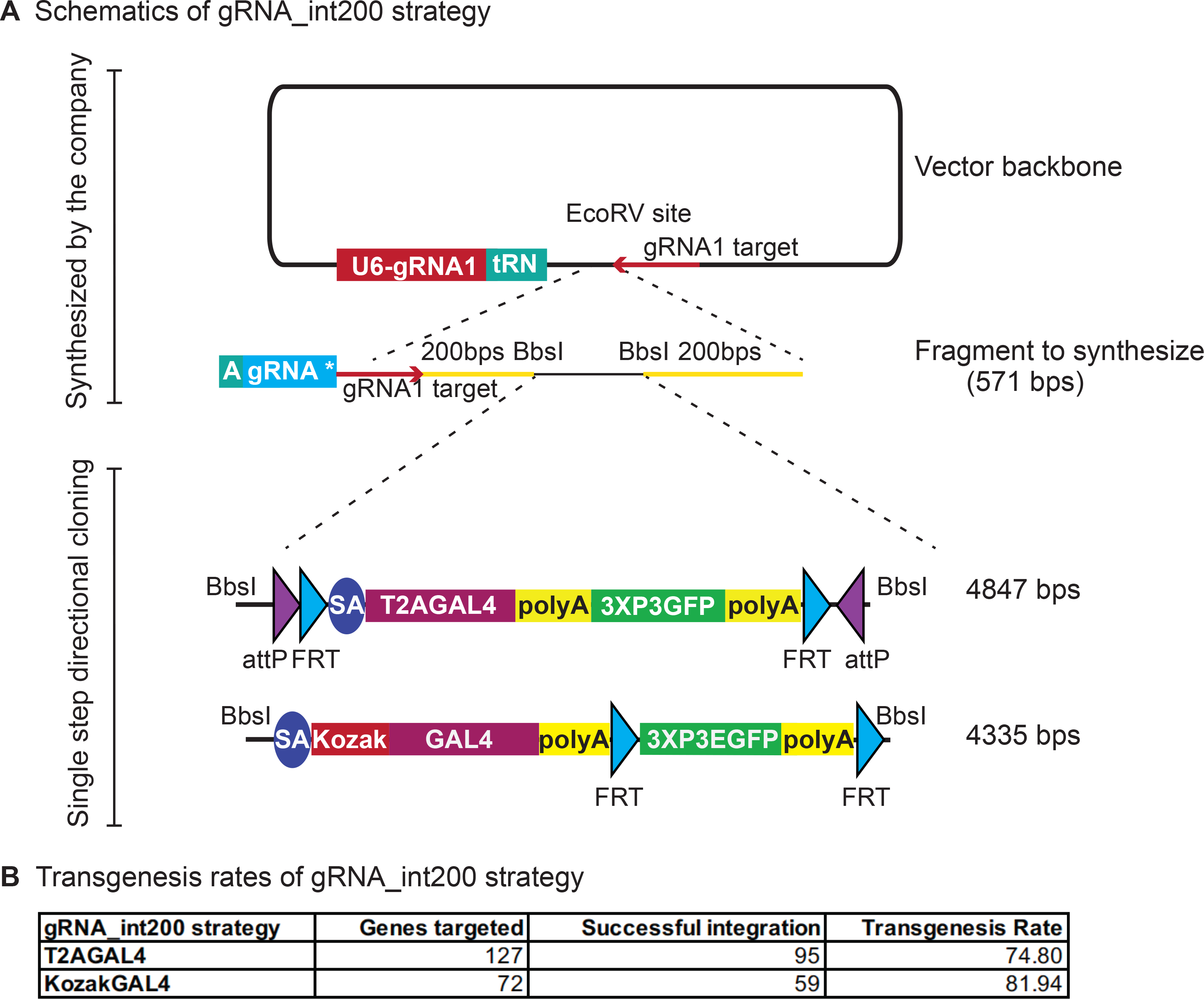
**gRNA_int200 strategy increases the transgenesis success rates. (**A) Schematics of the gRNA_int200 strategy. (B) Transgenesis data using gRNA_int200_T2AGAL4 or 2XgRNA_int200_KozakGAL4 strategies.

These data show that incorporating all the sgRNAs in the donor vector improves the transgenesis efficiency.

For the KozakGAL4 constructs, we inserted a second tRNA sequence after the first synthesized sgRNA and added the second gene specific sgRNA sequence to the synthesis reaction. We targeted 72 genes with 2XgRNA_int200_KozakGAL4 cassette and successfully inserted KozakGAL4 in 59 genes (∼82% success rate, Figure 4B; Supplemental Table 2). We tested whether genes for which the tagging failed using the int200_KozakGAL4 strategy (Figure 3) could be properly targeted with the 2XgRNA_int200_KozakGAL4 using the same gene specific sgRNAs and homology arms sequences and again observed that for 3 out of 4 tested genes, the 2XgRNA_int200_KozakGAL4 strategy was successful. In summary, the gRNA_int200 design increases transgenesis rate and streamlines the creation of T2AGAL4 and KozakGAL4 constructs by obviating the need to generate a separate sgRNA expression plasmid and ensuring co-delivery of all components for homologous recombination. In summary, the gRNA_int200 allows a 78% transgenesis success rate, or an additional 20% when compared to the int200 approach (increase from 65% to 78%).

### Use of 2XgRNA_int200 intermediate vectors for GFP tagging

We have previously shown that integrating a *SA-linker-EGFP-FlAsH-StrepII- 3xTEVcs-3xFLAG-linker-SD* (*SA-GFP-SD*) in coding introns of genes is an efficient approach to tag proteins with GFP (Venken *et al*., 2011; Nagarkar-Jaiswal *et al.,* 2015a; 2015b; 2017; Li-Kroeger *et al*., 2018). We typically generate these alleles through Recombinase Mediated Cassette Exchange of existing MiMIC SICs and we have shown that they are functional in 72% of tested genes (Nagarkar-Jaiswal *et al*., 2015a). A major factor that affects the functionality of the GFP protein trap is the insertion position. In cases where the artificial exon encoding for protein trap is inserted in a coding intron that bisects a predicted functional protein domain, the resulting protein trap is often not functional. Hence, another efficient approach to tag proteins encoded by genes that have no intron, small introns or no suitable MiMICS in any preselected position in the protein structure is highly desirable.

We tested the use of synthesized homology donor intermediate vectors to replace the coding sequence of genes without suitable coding introns with the gene coding sequence fused to GFP at different locations. We selected the *Wdr37* gene as it has a small intron (Kanca *et al*., 2019b). We amplified *Wdr37* sequences from the genome by PCR and used NEB HiFi DNA assembly to generate homology donor constructs where a sfGFP tag is integrated at the N terminus, C terminus or internally (Figure 5, Figure 5 Supplementary figure 1,2,3 for schematics of HiFi assembly). The 3XP3 DsRed flanked by PiggyBac transposase inverted repeats is integrated after the 3’UTR and serves as the transformation marker that can be excised precisely using the PiggyBac transposase (flyCRISPR.molbio.wisc.edu; Bier *et al*., 2018) (Figure 5A). The assembled sequences are subcloned in the synthesized homology donor intermediate. Injection of the homology donor plasmids in embryos expressing Cas9 in their germline resulted in positive transgenics in each case. Western blot of the resulting protein trap alleles using anti-GFP antibody detected bands at the expected length for the tagged protein in each case. However, the internally tagged allele is less abundant, underlining that the placement of sfGFP tag can affect protein stability (Figure 5B). Hence, the strategy to replace the whole coding region with a GFP tagged coding region allows tagging almost any gene in any position in the coding sequence.

**Figure 5.**
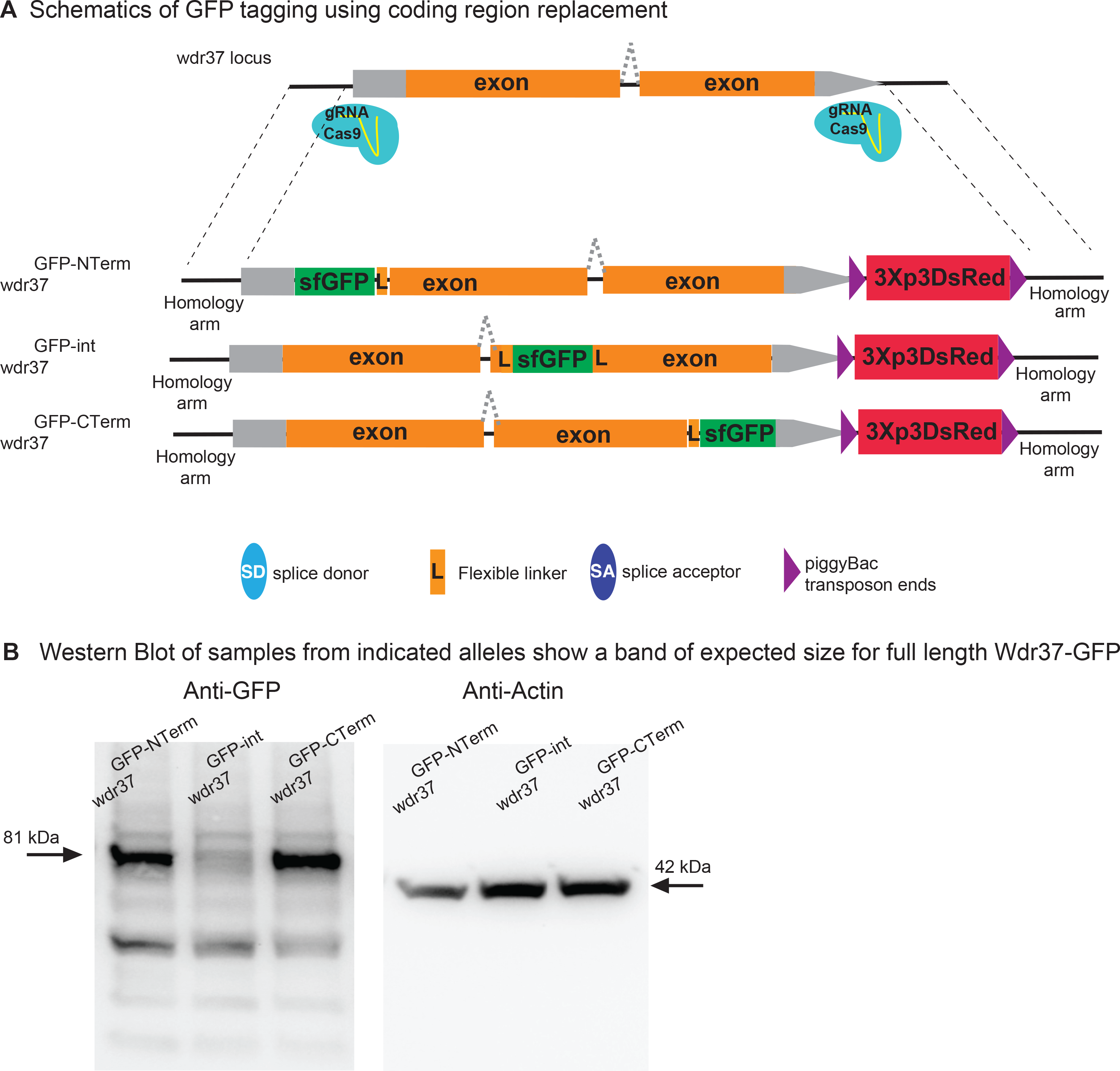
2XgRNA_int200 strategy can be used to tag any gene at any coding region to generate protein trap alleles. (A) Schematics of the targeting constructs to integrate sfGFP protein tag at an N-terminal, internal or C-terminal location in *Wdr37* gene locus. (B) Western blot analysis from adult flies show full length protein in all protein trap alleles with the arrow indicating the 81 kDa band that is the length predicted for the Wdr37 protein fused to sfGFP.

In summary, we developed a *KozakGAL4* strategy to target the genes that do not have a suitable intron and a set of novel custom vector backbones to facilitate homology donor construct production and increase transgenesis rate. The methods we developed are versatile and can be modified to generate GAL4 gene traps or GFP protein fusions of the targeted genes. Finally, the methodology we describe should be easy to implement in any other model organisms to facilitate generation of gene trap and protein trap alleles.

## Acknowledgements

We thank the Bloomington *Drosophila* Stock Center and FlyBase for fly stocks acquisition and distribution and the Developmental Studies Hybridoma Bank for antibodies. We thank Stephanie Mohr for critical reading of the manuscript. We thank Yuchun He, Ying Fang, Minhua Huang, Zhihua Wang, Yaping Yu, Junyan Fang, Ruifang Zhang, and Lily Wang for generating and maintaining MiMIC/CRIMIC *T2A-GAL4* fly stocks. Confocal microscopy was performed in the BCM IDDRC Neurovisualization Core, supported by the NICHD (U54HD083092). HJB received support from NIH R01GM067858, R24 OD031447, R24OD022005, U54NS093793 HJB and the Huffington Foundation. OK receive support from NIH R24 OD031447. JZ, YHu, and NP receive support from NIGMS (GM067761 and GM084947). NP is an investigator of the Howard Hughes Medical Institute. RWL was supported by the Carnegie Institution for Science.

## Materials and Methods

### Generation of homology donor constructs

Templates for ordering the int200 and gRNA_int200 constructs can be found in Supplementary Materials and Methods. Homology donor intermediate vectors were ordered for production from Genewiz (“ValueGene” option) in pUC57 Kan_gw_OK (for int200 strategy) or pUC57 Kan_gw_OK2 (for gRNA_int200 strategy) vector backbone at 4 μg production scale. The lyophilized vectors were resuspended in 53 μl of ddH2O. 1 ul was used for Golden Gate assembly with 290 ng of pM37 vector of reading frame phase corresponding to the targeted intron (for T2GAL4, Lee *et al*., 2018) or 265 ng of pM37_KozakGAL4 vector (for KozakGAL4). The Golden Gate Assembly reaction was set in 200µl PCR tubes (ThermoScientific AB2000) with 2.5 µl 10X T4 DNA ligase buffer (NEB B0202S), 0.5 µl T4 DNA ligase (NEB M0202L), 1 µl restriction enzyme (BbsI_HF or BsaI_HFv2 NEB R3559L and R3733L respectively), 1 µl of SIC (pM37_T2AGAL4 or pM37_KozakGAL4 at 290 ng/ µl or 265 ng/ µl respectively), 19 µl of dH2O and 1 µl of homology donor construct. For cloning multiple constructs in parallel, master mixes were prepared including all the components except for the homology donor intermediate vector. The reactions were incubated in a Thermocycler (cycle 30 times between 37°C 5 minutes, 16°C 5 minutes, then 65°C 20 minutes, 8°C hold). An additional digestion step was done to remove self ligating plasmid backbones by adding 19.5 µl dH2O, 5 µl 10X CutSmart buffer (NEB B7204S) and 0.5 µl BbsI or BsaI_HFv2 (the enzyme used for the cloning reaction). The reaction product was transformed in DH5⍺ competent cells and plated on Kanamycin+ LB plates.

### Fly injections

Int200-T2AGAL4 and int200-KozakGAL4 constructs were injected at 250 ng/µl along with 100ng/µl gene specific gRNA(s) cloned in pCFD3 or pCFD5 respectively (Port *et al*. 2014; Port and Bullock, 2016). Injections were performed as described in (Lee *et al*., 2018). 400-600 embryos from *y^1^w*; iso18; attP2(y+){nos-Cas9(v+)}* for genes on the 2^nd^ or 4^th^ chromosome and *y^1^w* iso6*;; *attP2(y+){nos-Cas9(v+)}* for genes on the X chromosome and *y^1^w*; attP40(y+){nos-Cas9(v+)}; iso5* (Kondo and Ueda, 2013) for genes on the 3^rd^ chromosome per genotype were injected. Whole genome sequencing BAM files of isogenized lines can be found at: https://zenodo.org/record/1341241. Resulting G0 males and females were crossed individually to *y^1^ w\** flies as single fly crosses for *3XP3-EGFP* detection. Positive lines were balanced, and stocks were established. Up to 5 independent lines were generated per construct per gene. The list of generated alleles can be found on Supplementary table 2. The sequences of homology arms and sgRNA(s) as well as the results of PCR validation and imaging on third instar larval brain are available at http://flypush.imgen.bcm.tmc.edu/pscreen/crimic/crimic.php. The stocks are deposited in the Bloomington Drosophila Stock Center (BDSC) on a regular basis. The stocks are available from the Bellen lab until they are deposited and established in the BDSC.

### PCR validation

PCR primers that flank the integration site were designed for each targeted gene. These primers were used in combination with insert-specific primers that bind 5’ of the inserted cassette in reverse orientation and 3’ of the insert in forward orientation (pointing outwards from the insert cassette, Primer sequences can be found in the supplementary material). 200-800 nt amplicons were amplified from genomic DNA from individual insertion lines through single fly PCR (Gloor et al., 1993) using OneTaq PCR master mix (NEB #M0271L). PCR conditions were 95°C for 30 seconds, 95°C 30 seconds, 58°C 30 seconds, 68°C 1 minute for 34 cycles and 68°C 5 minutes.

### Confocal imaging of transgenic larval brains

Dissection and imaging were performed following the protocols in (Lee *et al*., 2018). In brief, fluorescence-positive 3^rd^ instar larvae were collected in 1x PBS solution and then cut in half and inverted to expose the brain. Brains were transferred into 1.5mL centrifuge tubes and fixed in 4% PFA in 1xPBS buffer for 20 minutes. Brains were then washed for 10 minutes three times in 0.2% PBST. Finally, samples were mounted on glass slides with 8µL of VectaShield (VectorLabs #H-1000) and imaged at 20x zoom with a Nikon W1 dual laser spinning-disc confocal microscope.

### Analysis of single cell sequencing data

To identify genes with expression profiles that overlap with expression of genes replaced with *KozakGAL4* sequences, we queried the data from third instar larval CNS scRNAseq data described in Ravenscroft *et al*. (2020). The data (http://scope.aertslab.org/#/Larval_Brain/*/welcome) were imported into Seurat (version 4.0.1). Cells expressing the selected genes, for which the KozakGAL4 allele was generated (e.g *CG3770*, *CG10939*, *CG10947* and *CG15093*), were identified using WhichCells function and genes enriched in these cells were identified using FindMarkers with default parameters. A list of the top 10 genes that were minimally expressed outside the expression domain of genes with KozakGAL4 alleles was generated. We then selected genes from the list for which T2AGAL4 were generated and compared the expression profiles using available images.

### Western blots

Flies were homogenized using Cell Lysis Buffer (25 mM Tris-HCl pH 7.5, 100 mM NaCl, 1 mM EDTA, 1% Triton-X 100, 1X liquid protease inhibitor (Gen DEPOT), 0.1 M DTT). The supernatant was collected after centrifugation at 13, 000 rpm for 10 min at 4°C (Eppendorf 5424R with rotor Eppendorf FA-45-24-11). The supernatant was mixed with Laemmli Buffer containing β-mercaptoethanol and heated at 95°C for 10 min. Subsequently, the samples were loaded in 4–20% gradient polyacrylamide gels (Bio-Rad Mini-PROTEAN^®^ TGX™). Following electrophoresis, proteins were transferred onto a polyvinylidene difluoride membrane (Immobilon, Sigma). The membrane was blocked using skimmed milk and treated with the primary antibody for overnight. The following antibodies were used in the present study: rabbit anti-GFP (1:1000) (Thermo Fisher Scientific, #A-11122), mouse anti-Actin (1:5000) (EMD Millipore, #MAB1501). Horseradish peroxidase-conjugated secondary antibody was used to detect the respective primary antibody. Blots were imaged on a Bio-Rad ChemiDocMP.

### Cloning of *Wdr37-KIGFP* constructs

The fragments that position the GFP tag to the selected sites were PCR amplified from genomic DNA, sfGFP was amplified from pBS_SA_sfGFP_SD (Kanca *et al*., 2019a) and scarless DSred from pScarlessHD-DsRed (pScarlessHD-DsRed was a gift from Kate O’Connor-Giles (Addgene plasmid # 64703 ; http://n2t.net/addgene:64703 ; RRID:Addgene_64703). The fragments were used together with homology donor intermediate for *Wdr37* gene used for generating *KozakGAL4* allele (CR70111) cut with BbsI-HF to assemble NEB-HiFi DNA assembly following manufacturer’s instructions. Schematics of HiFi assembly can be found in Figure 5 Supplementary figures 1, 2 and 3.

---

**Supplementary Figure 1.**
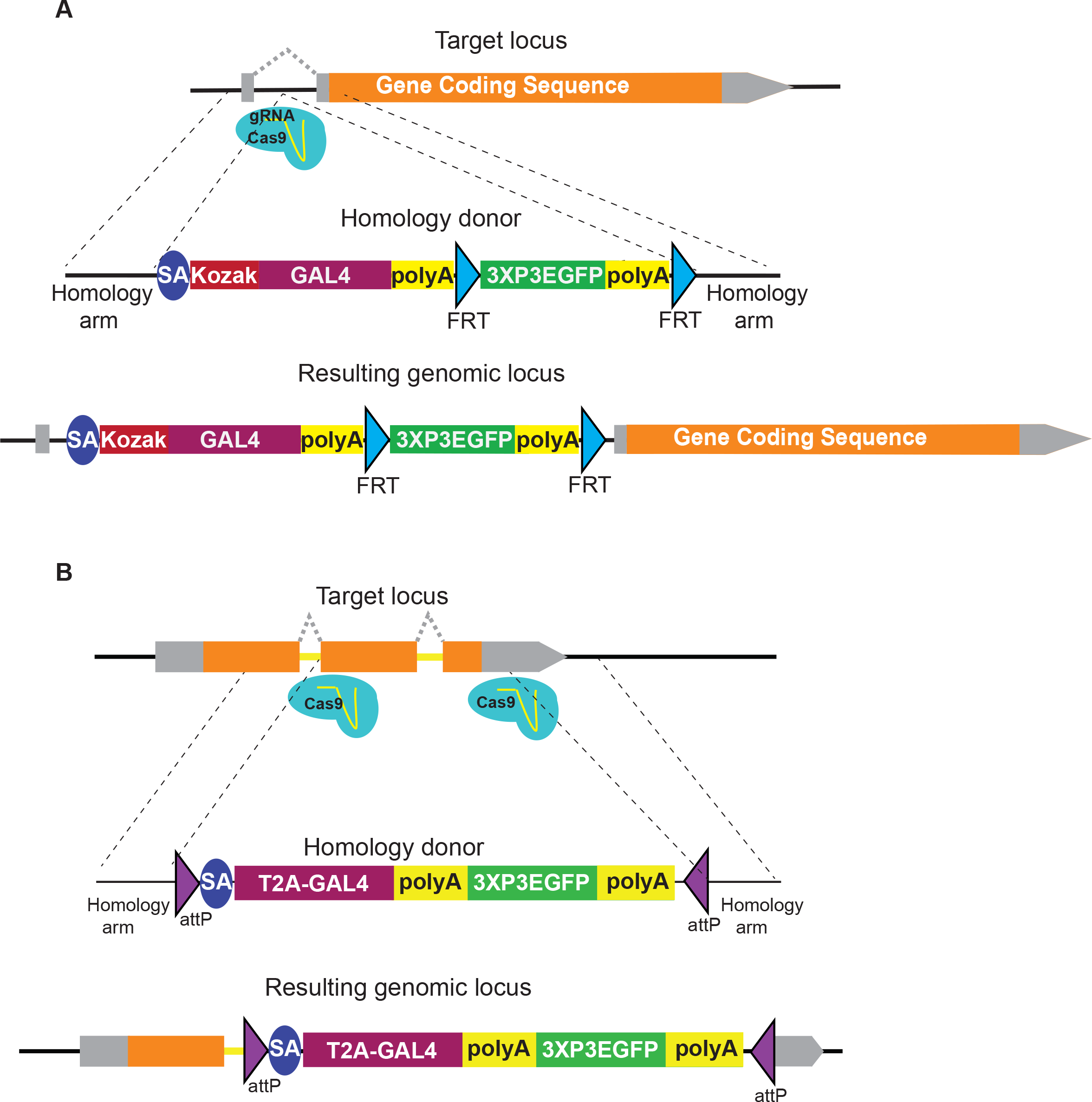
Alternative strategies to generate gene trap alleles in genes without suitable introns. For genes that cannot be targeted by artificial exon strategies and where suitable sgRNAs could not be found in the 5’UTR an artificial exon with SA_KozakGAL4 can be inserted in an intron in the 5’UTR (A) or in a short intron by deleting the exons following the intron (B).

**Supplementary Figure 2.**
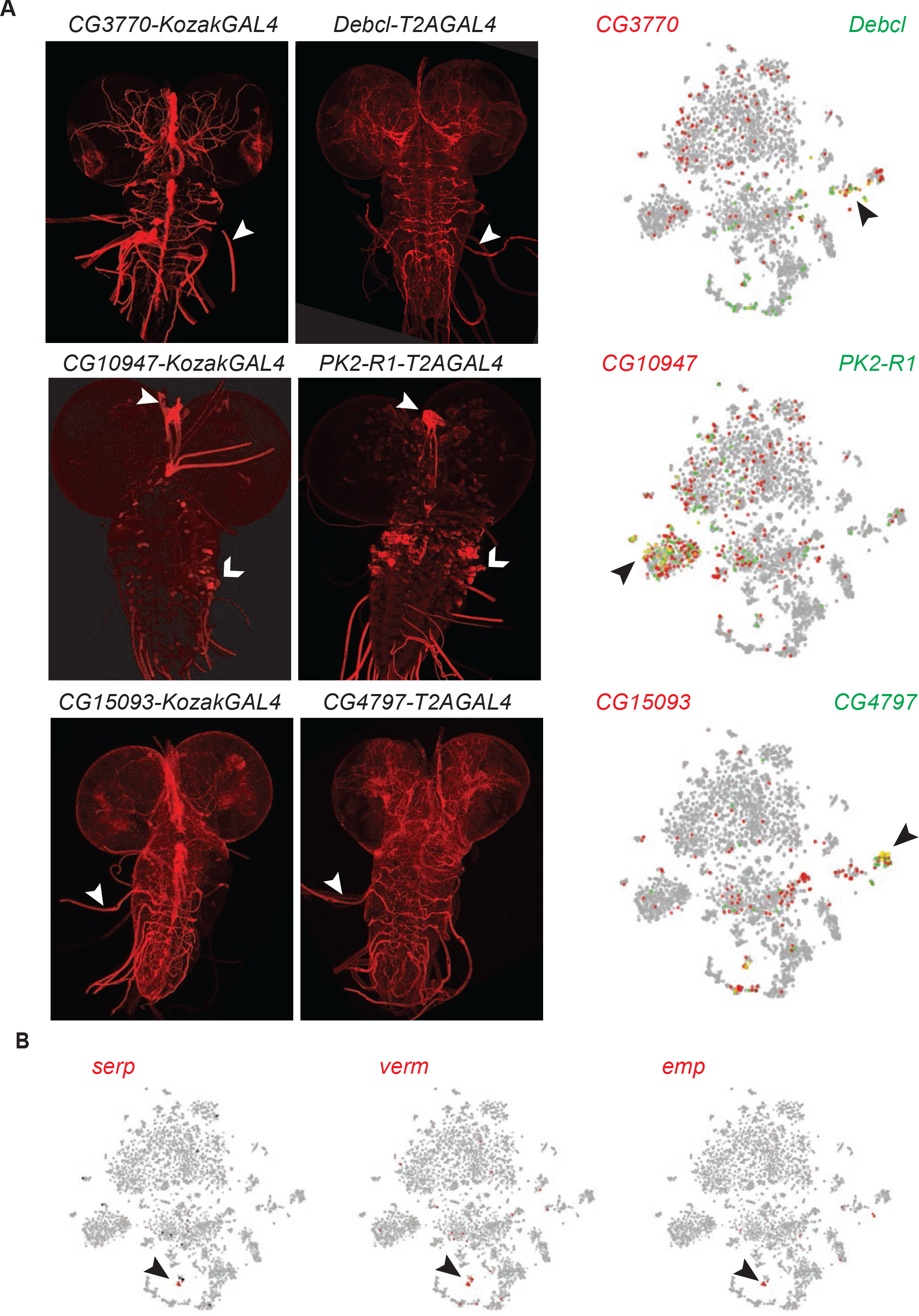
Identification of genes expressed in similar cells to the *KozakGAL4* alleles expressed in restricted patterns. (A) The single cell sequencing data from Ravenscroft *et al*. 2020 with the cells expressing the gene targeted by KozakGAL4 marked by red circles and cells expressing the gene targeted with T2AGAL4 allele marked by green circles. The imaging results of reporter expression generated with *KozakGAL4* alleles were compared to the expression patterns of genes that are expressed in similar cells by analysis of single cell sequencing data imaged using *T2AGAL4* alleles. Images are taken by crossing the *GAL4* alleles with *UAS-CD8mCherry.* Arrowheads show the regions with the most overlap. (B) Cluster of trachea markers in the scRNA data from Ravenscroft *et al*. 2020.

**Figure 5 Supplementary Figure 1.**
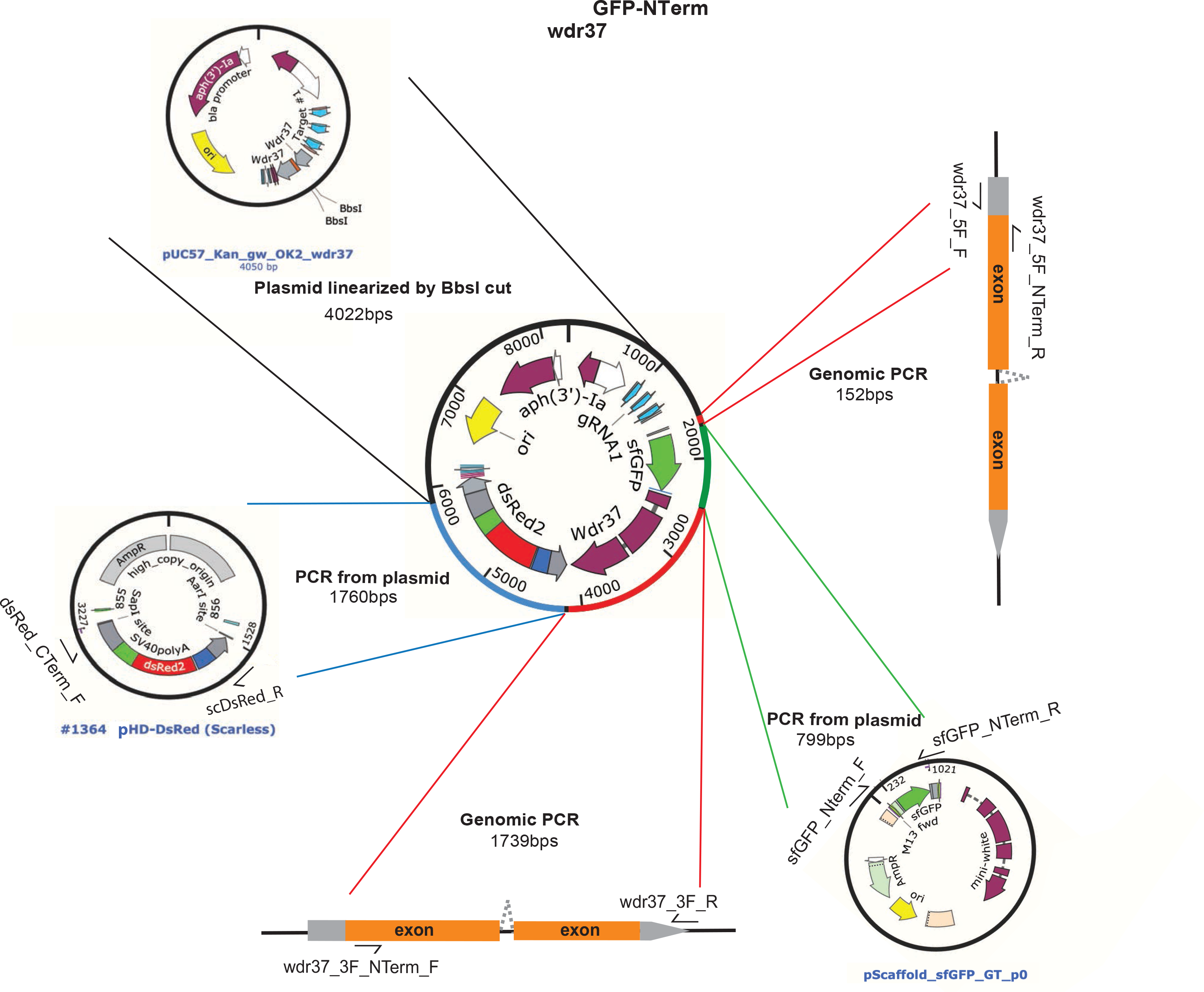
wdr37 GFP-NTerm protein trap allele donor constructs fragments schematics.

**Figure 5 Supplementary Figure 2.**
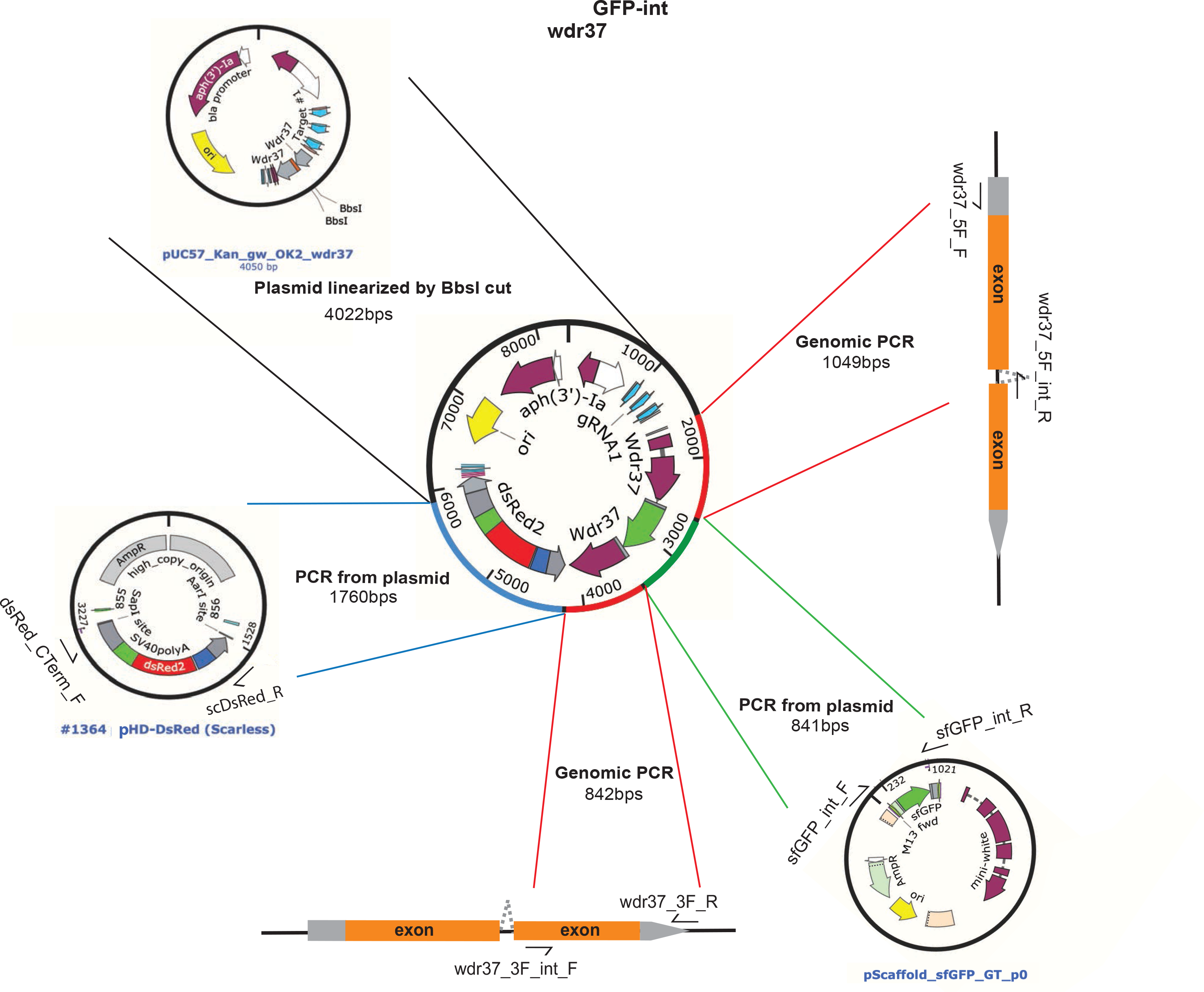
wdr37 GFP-NTerm protein trap allele donor constructs fragments schematics.

**Figure 5 Supplementary Figure 3.**
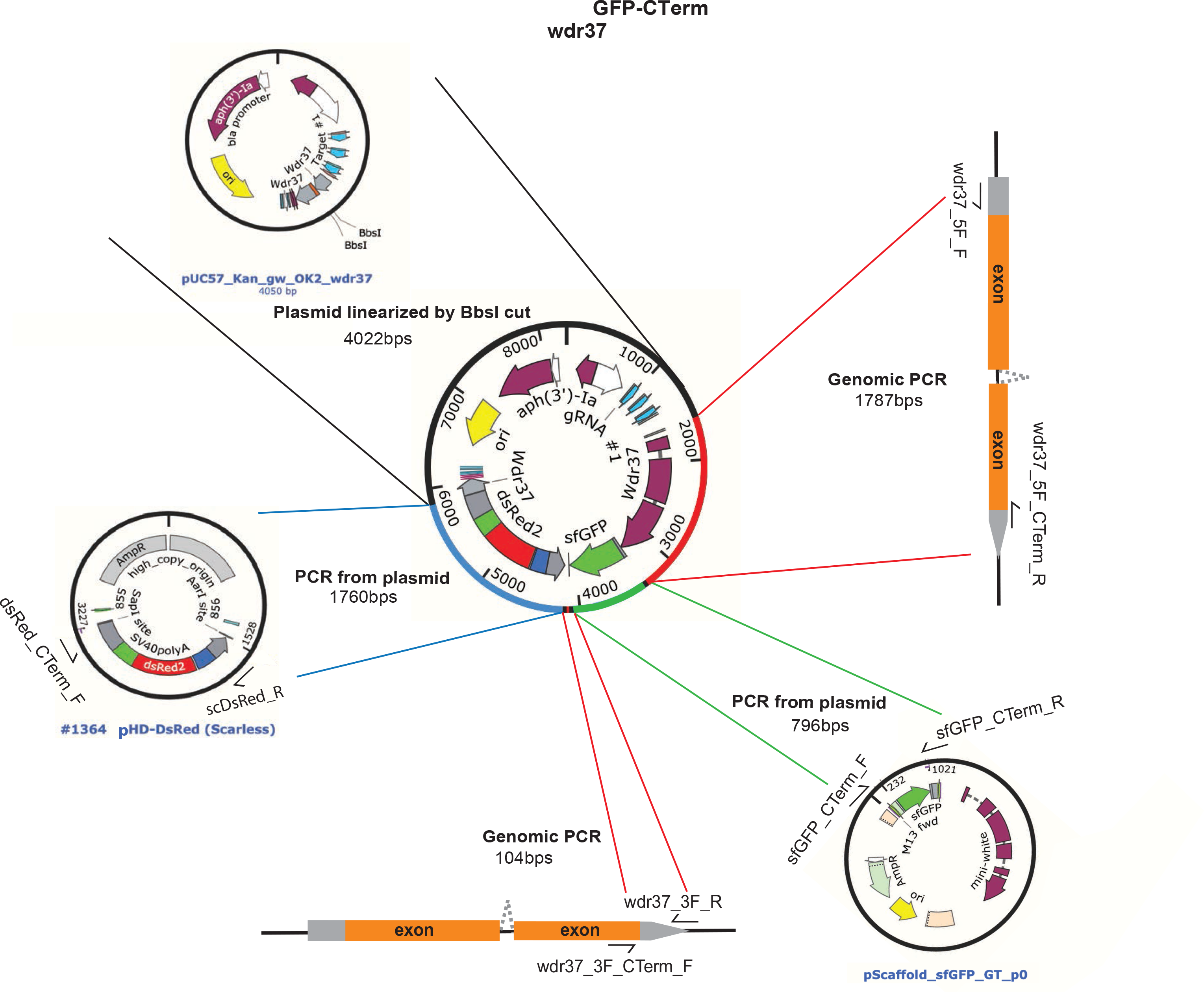
wdr37 GFP-CTerm protein trap allele donor constructs fragments schematics.

**Supplementary Table 1.**
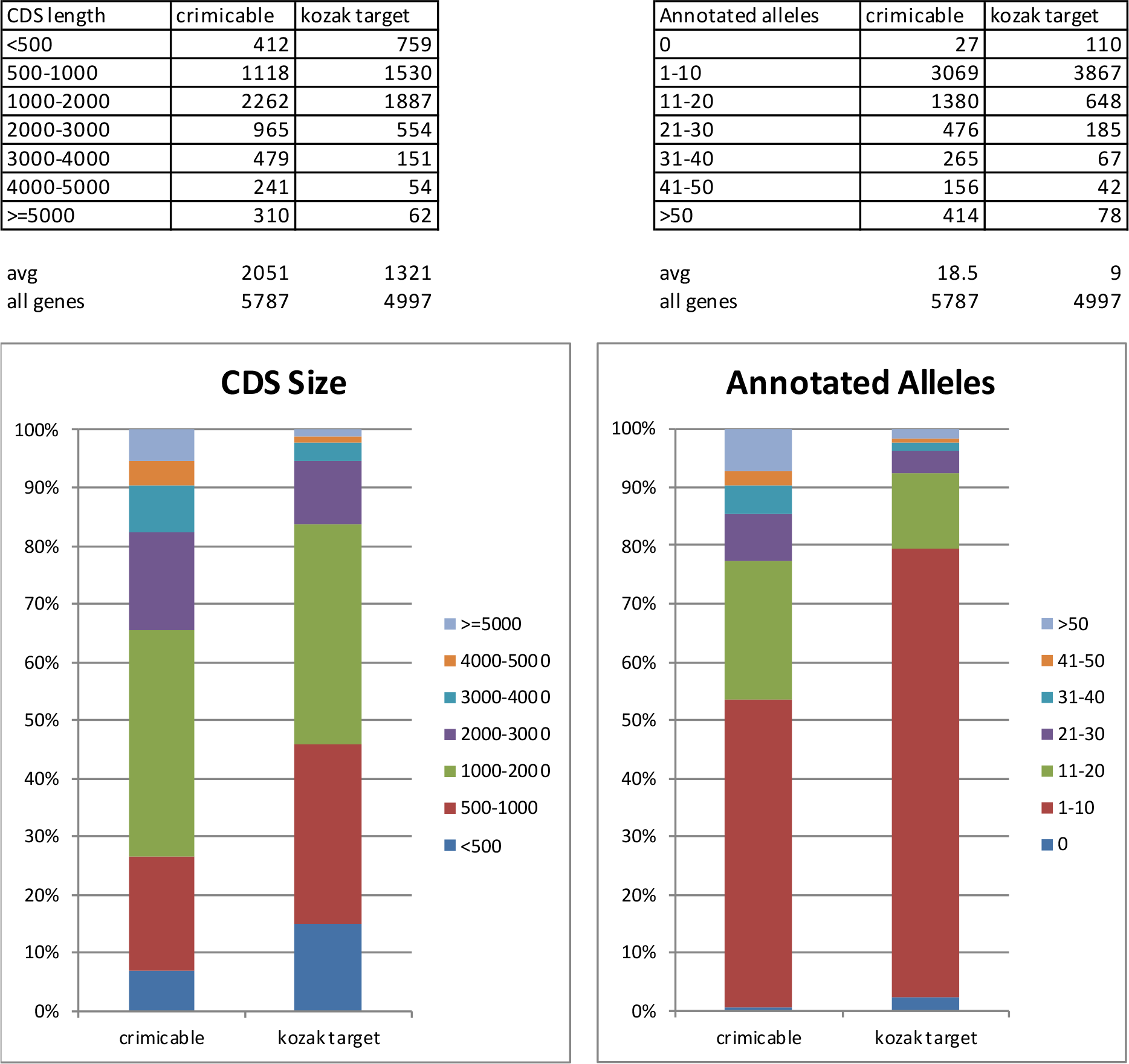
Analysis of Drosophila genome for the presence of suitable introns.

**Supplementary Table 2.**
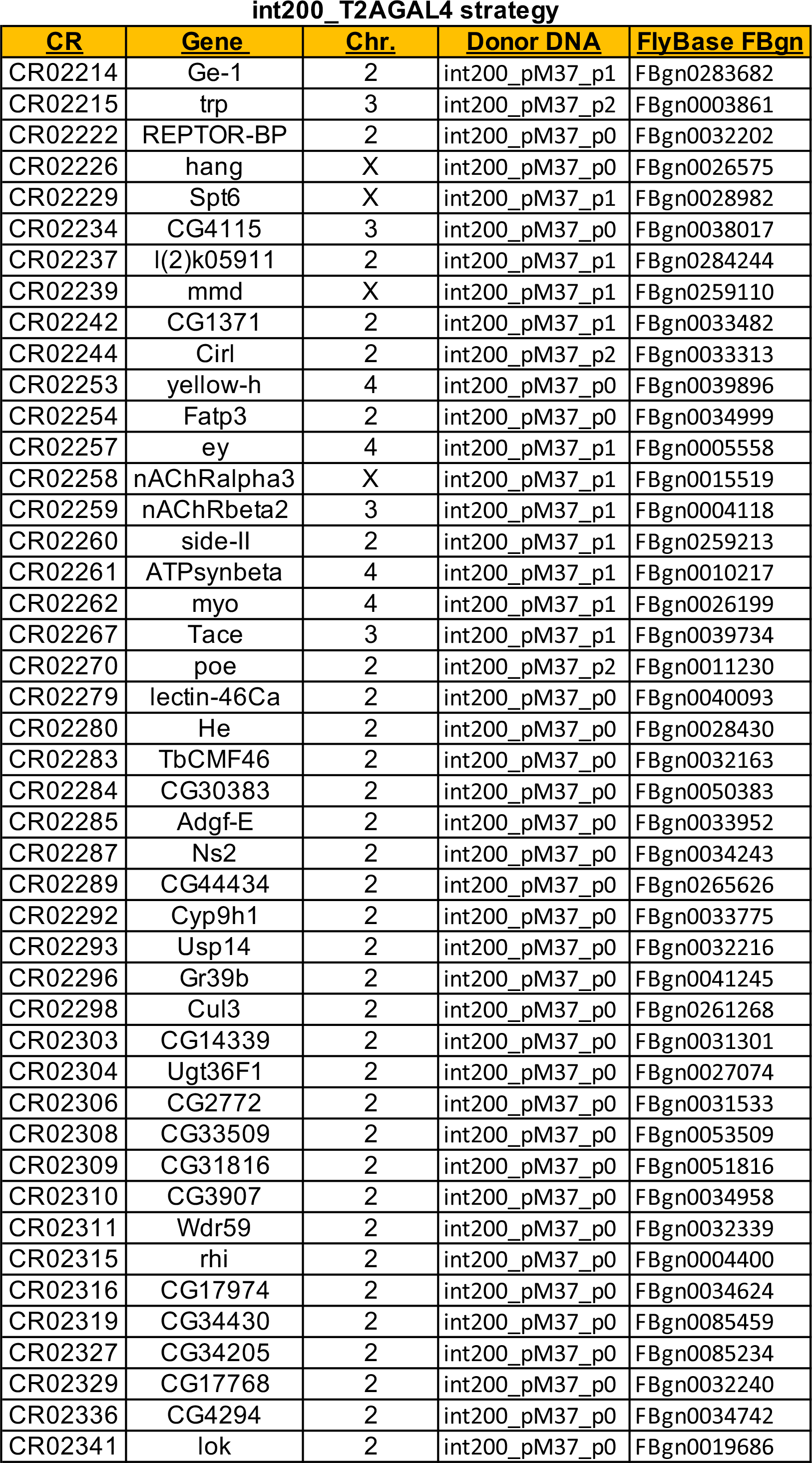
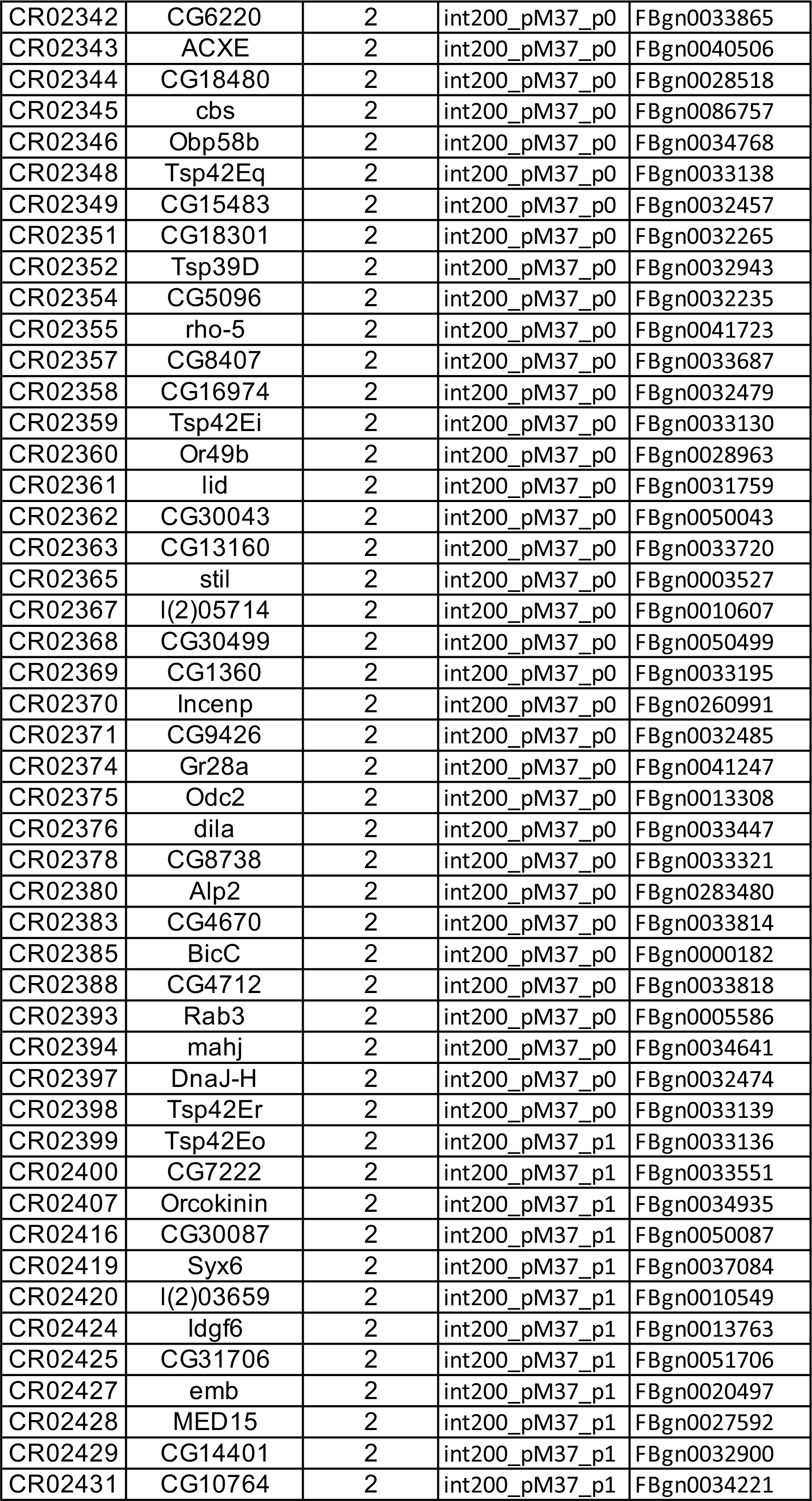
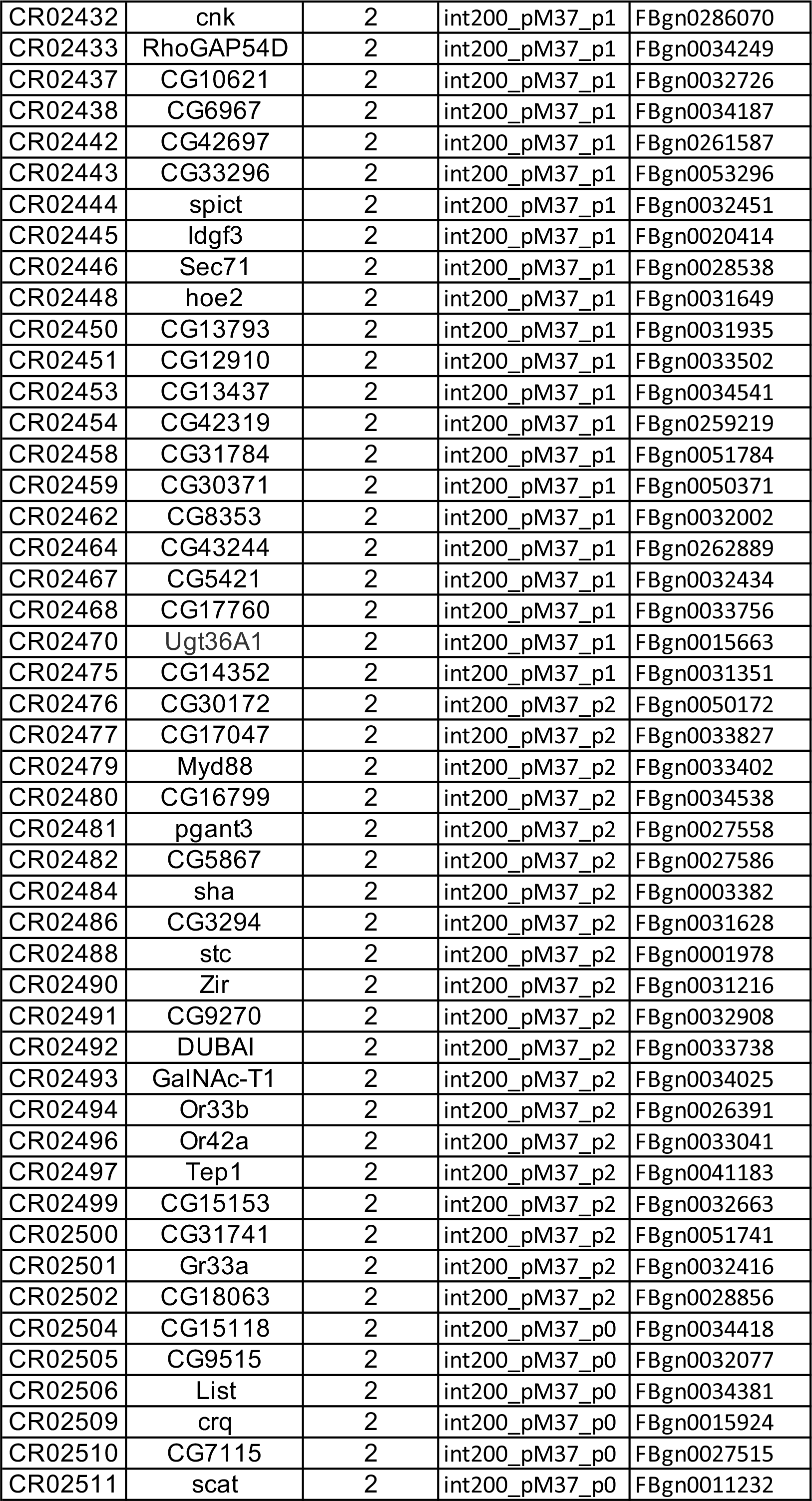
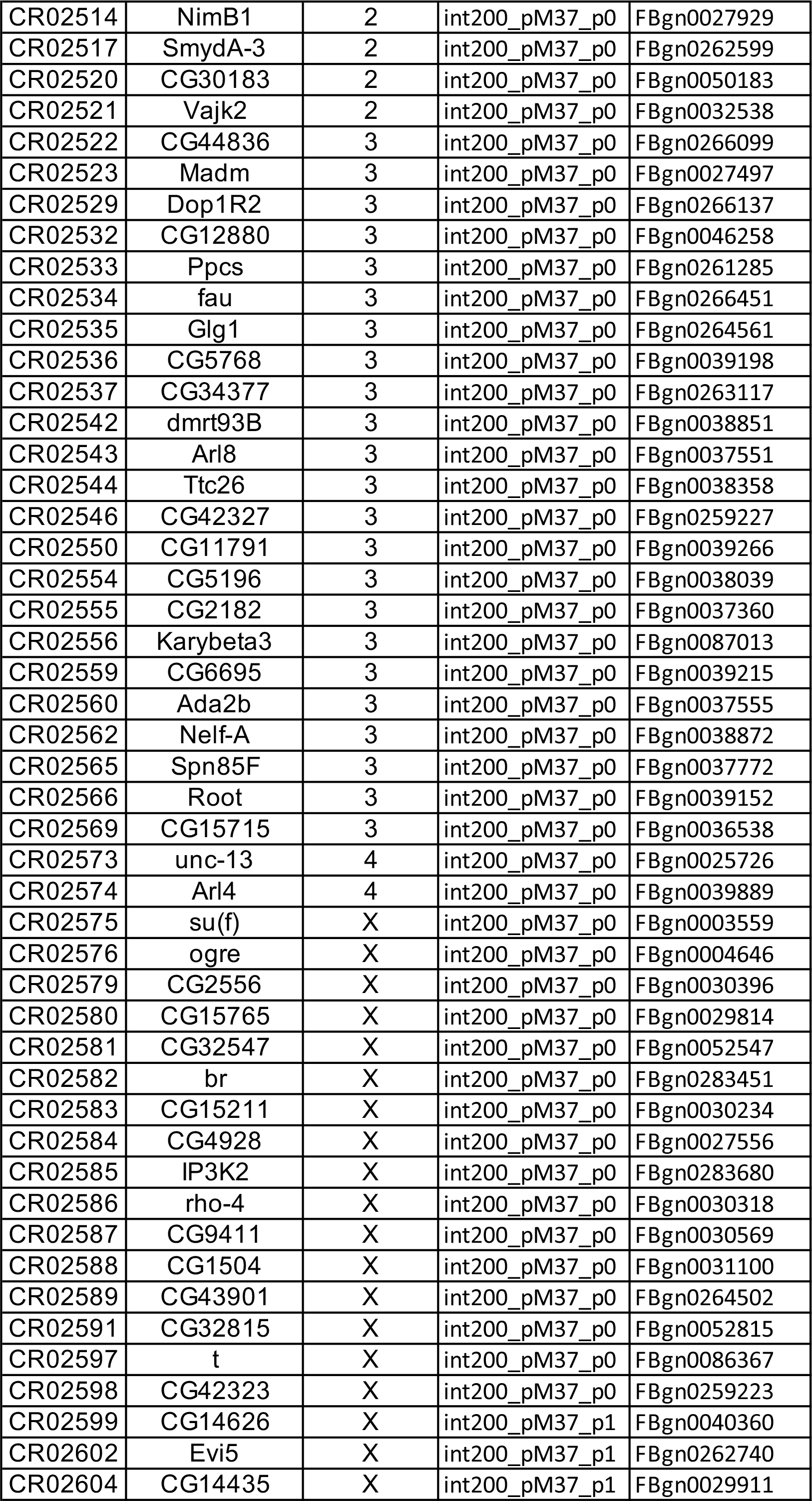
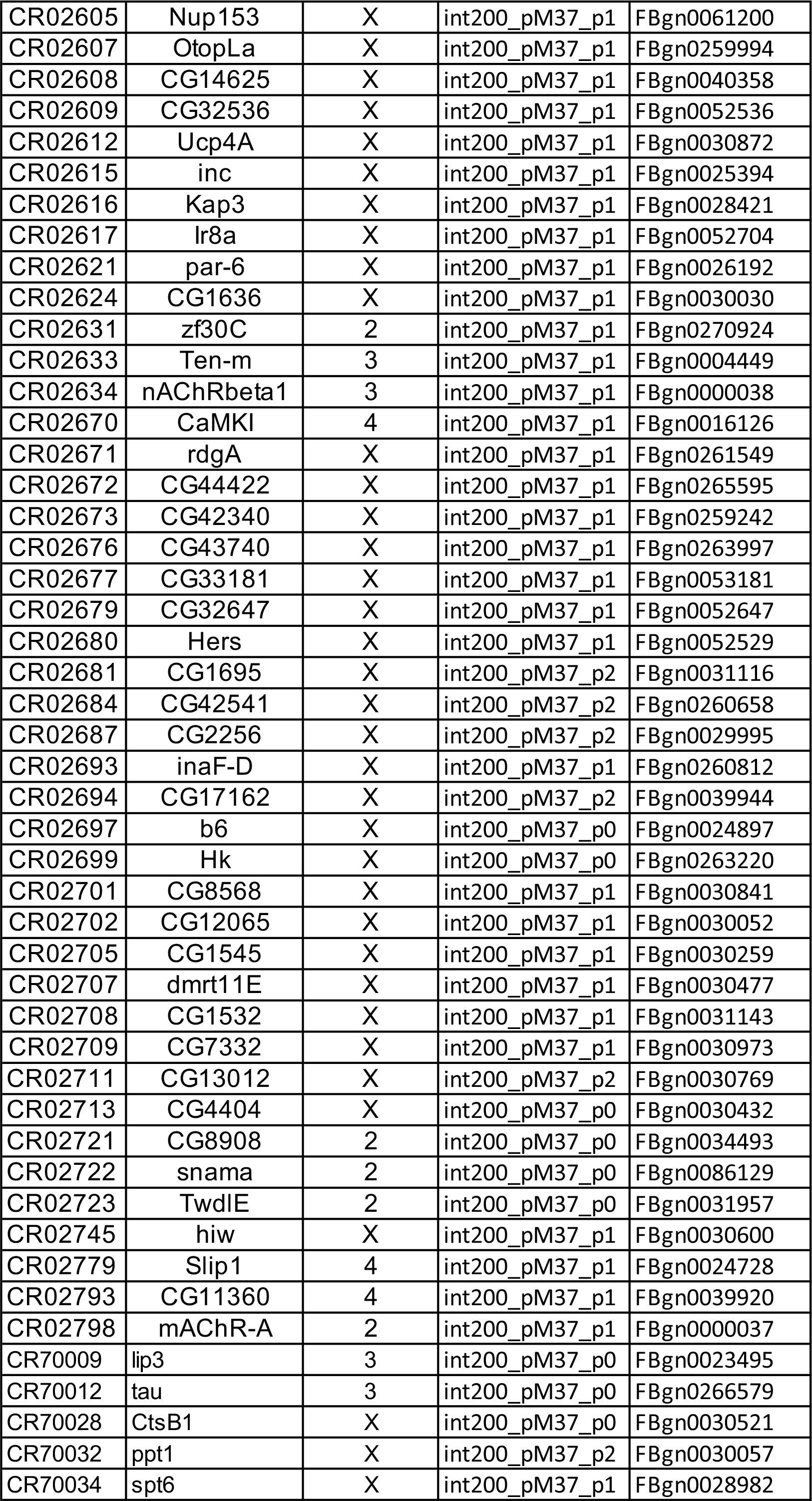
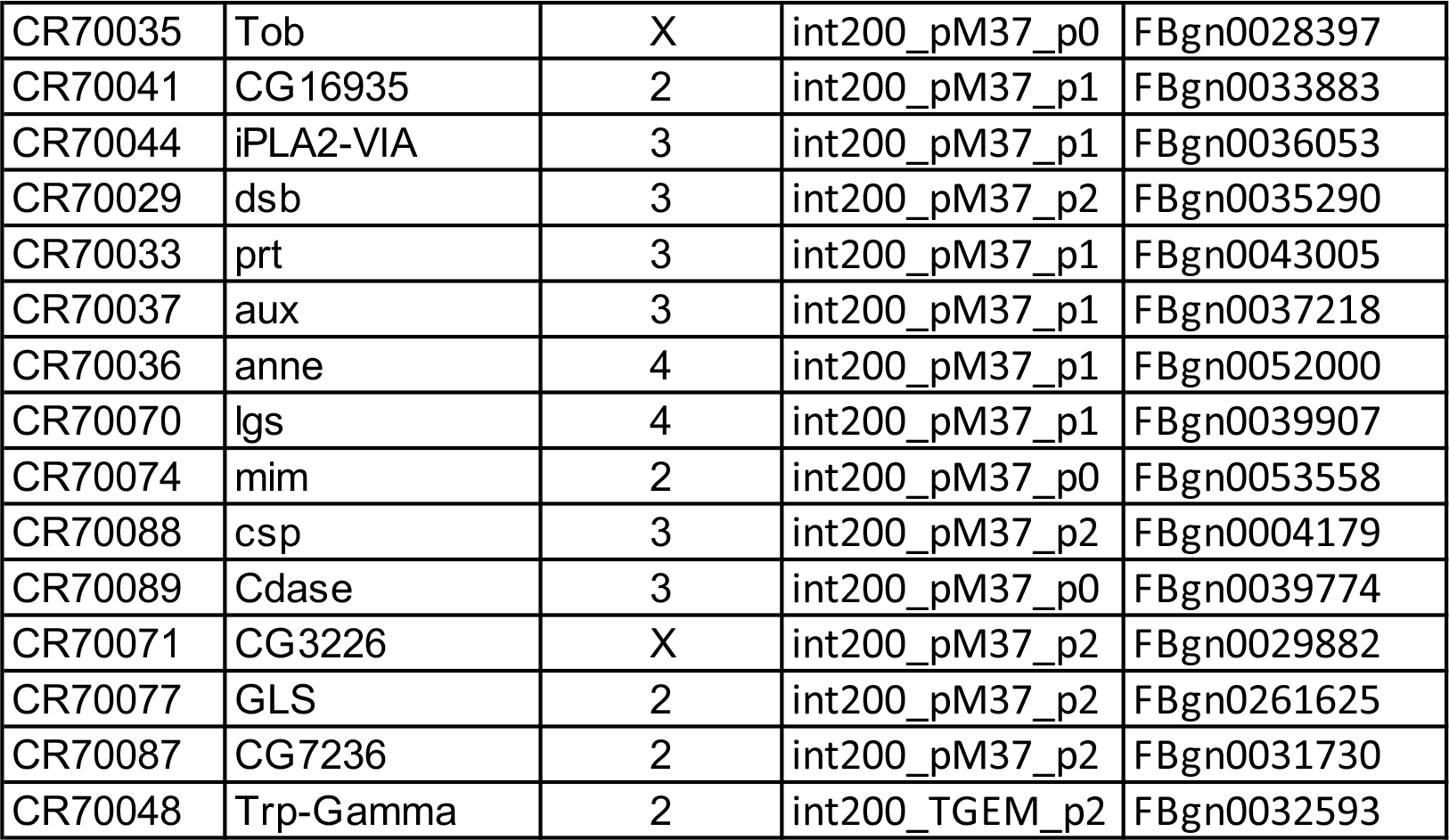
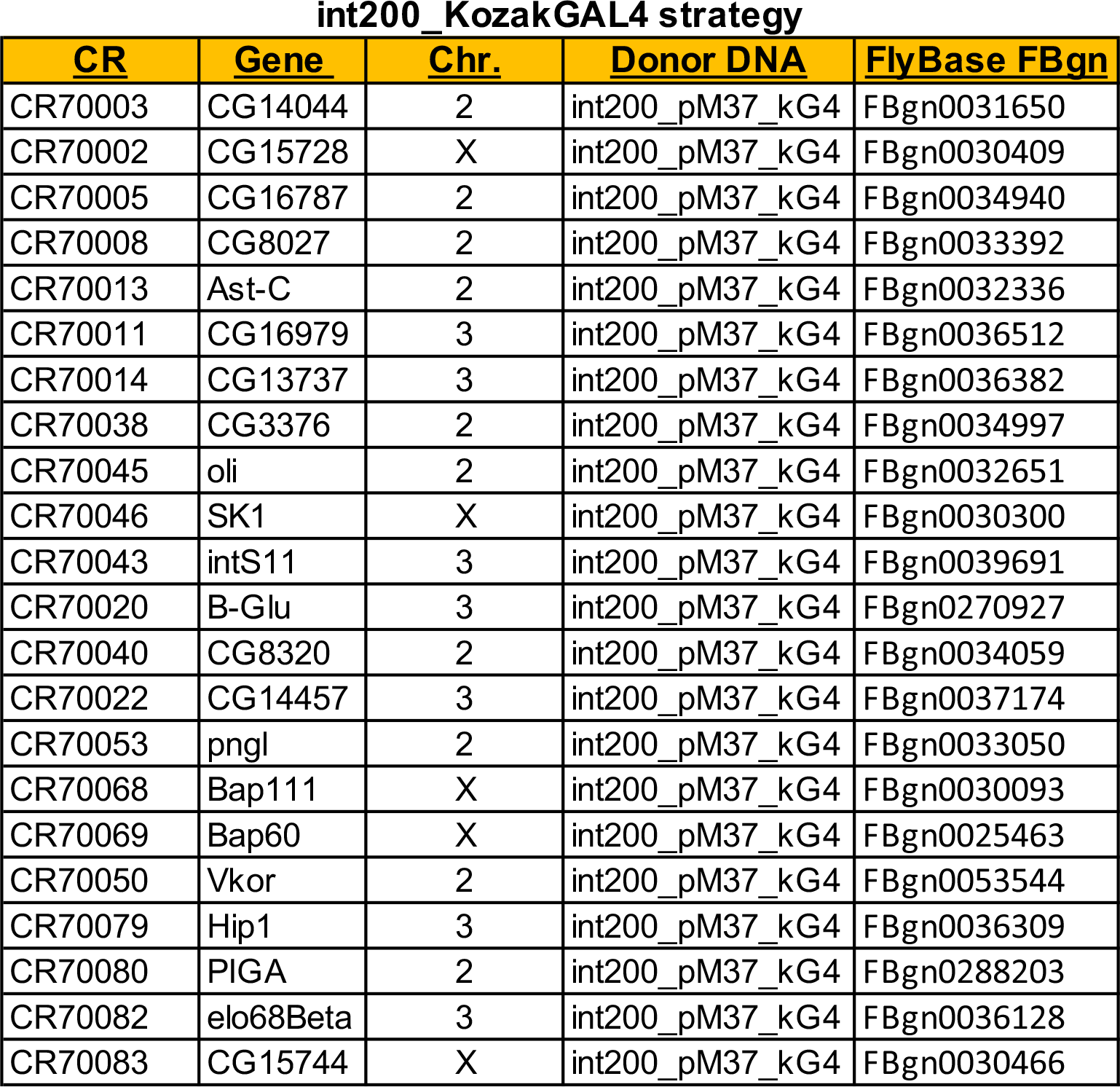
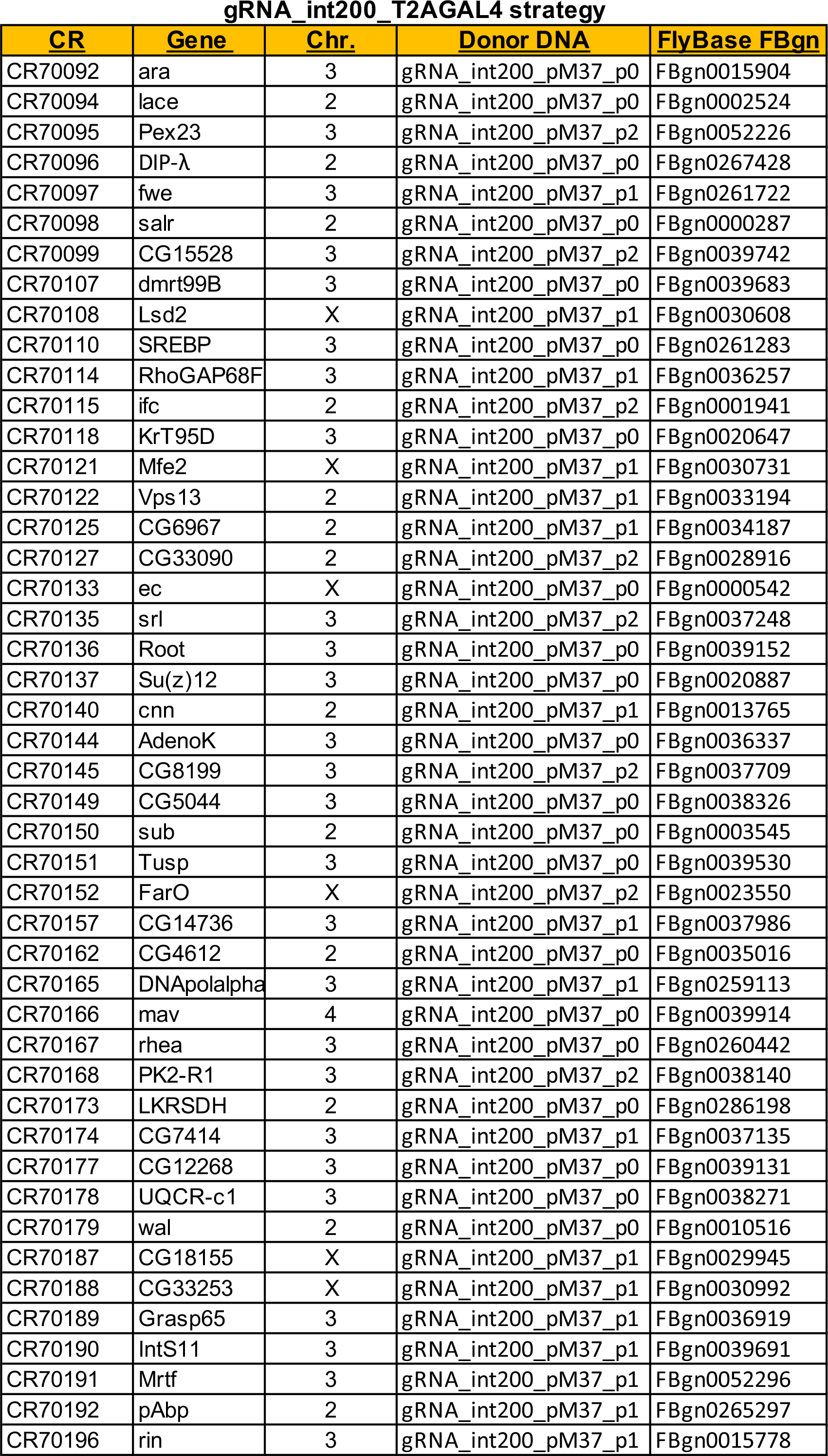
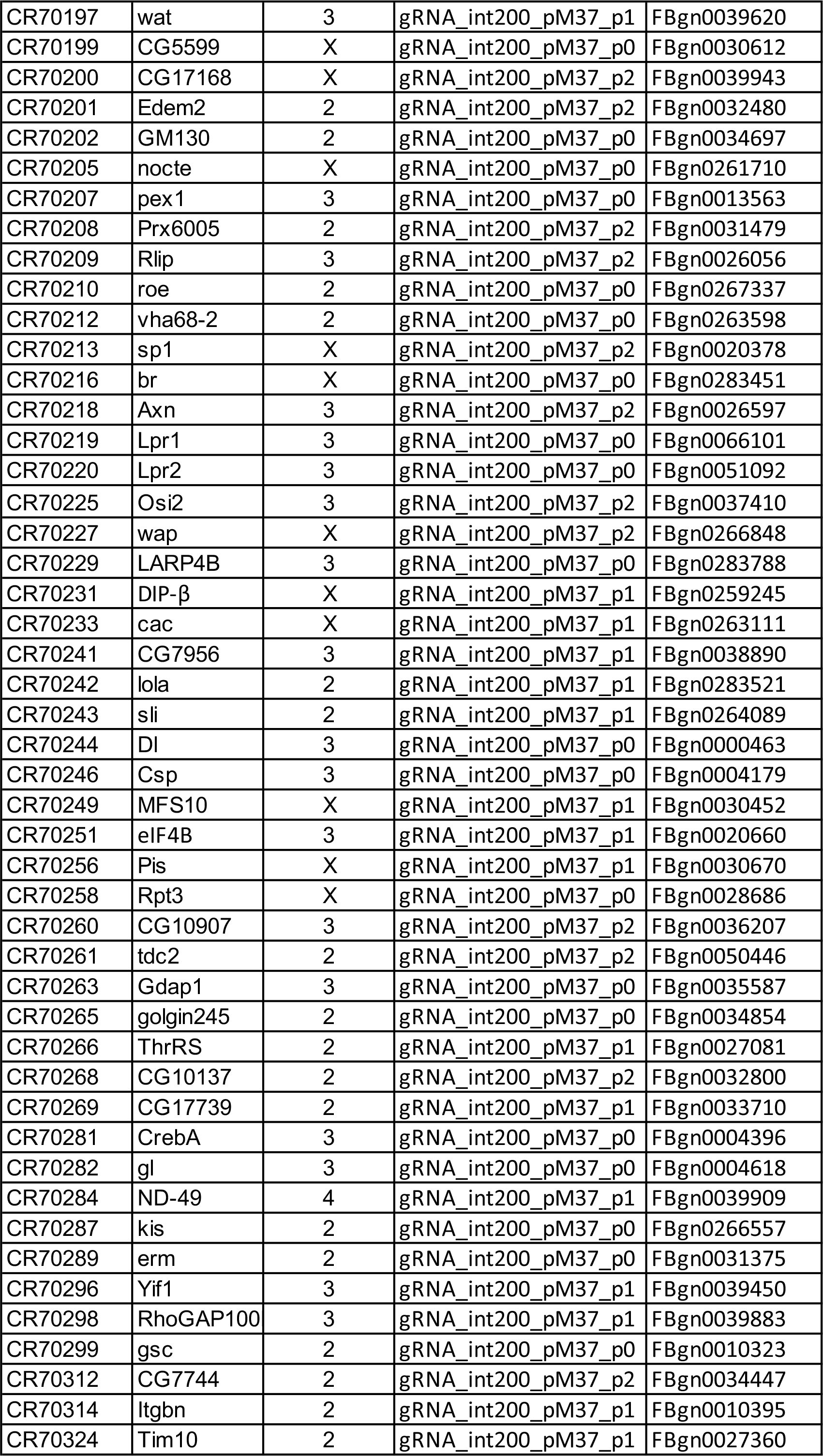
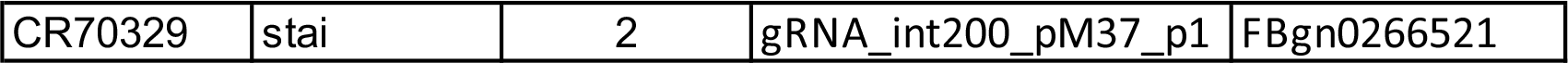
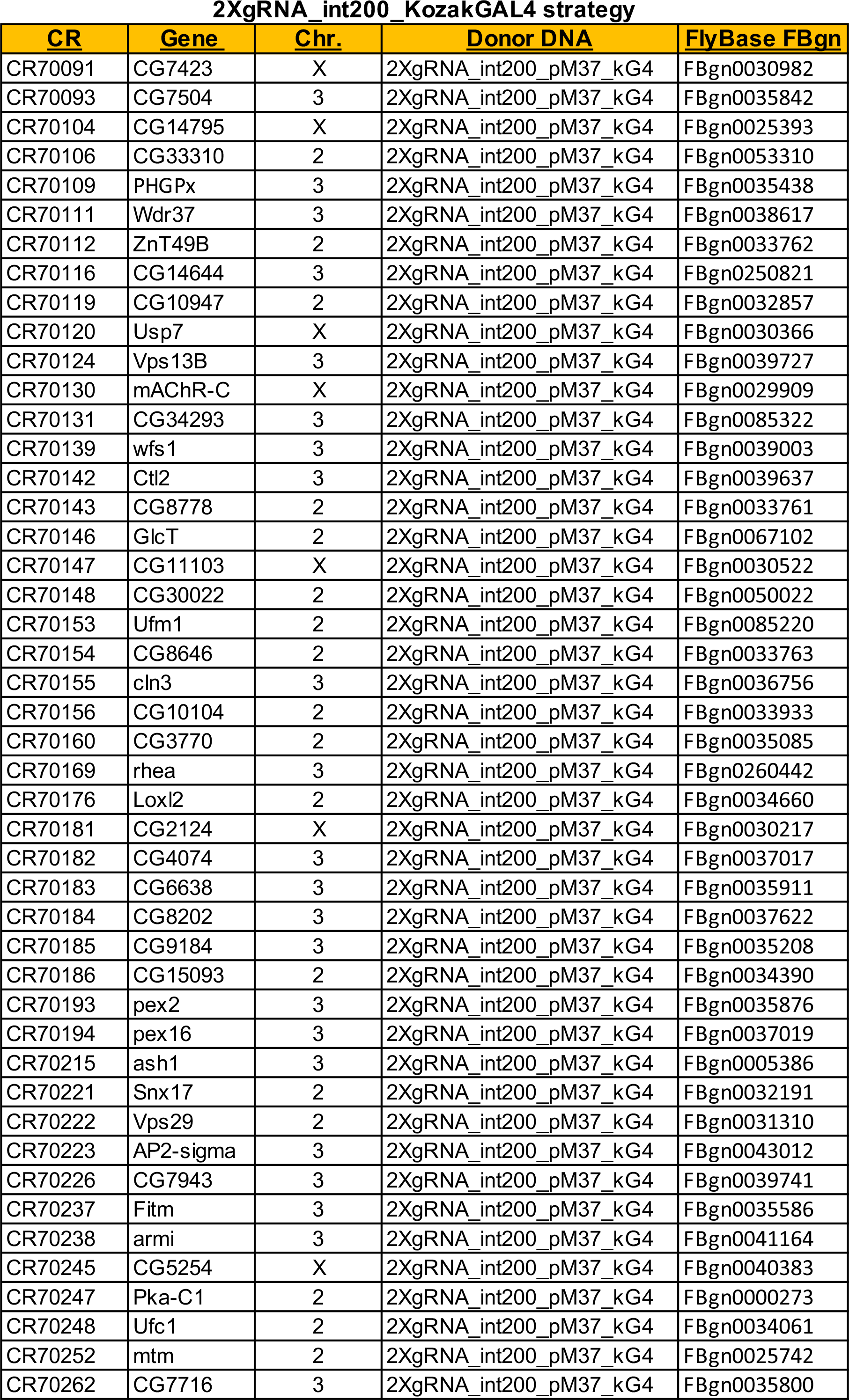
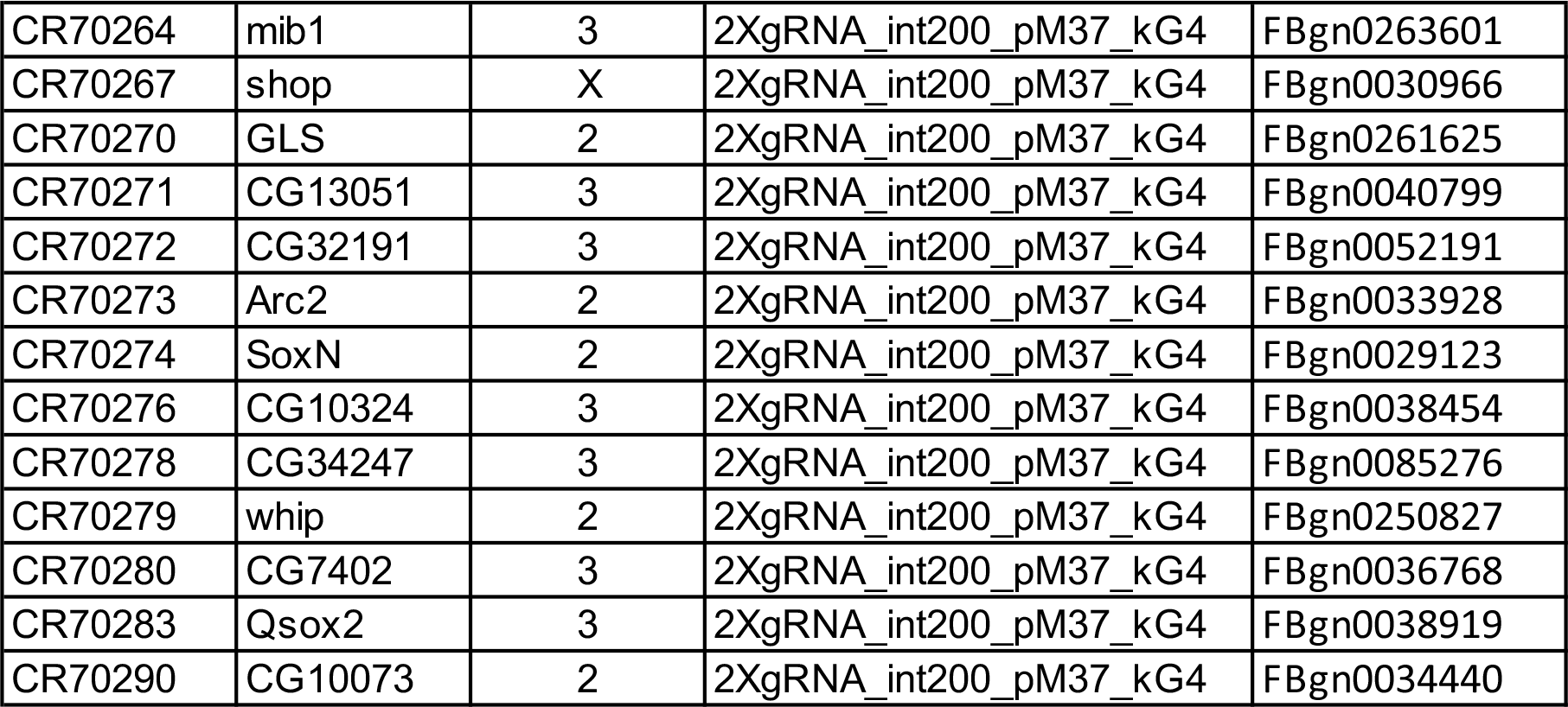
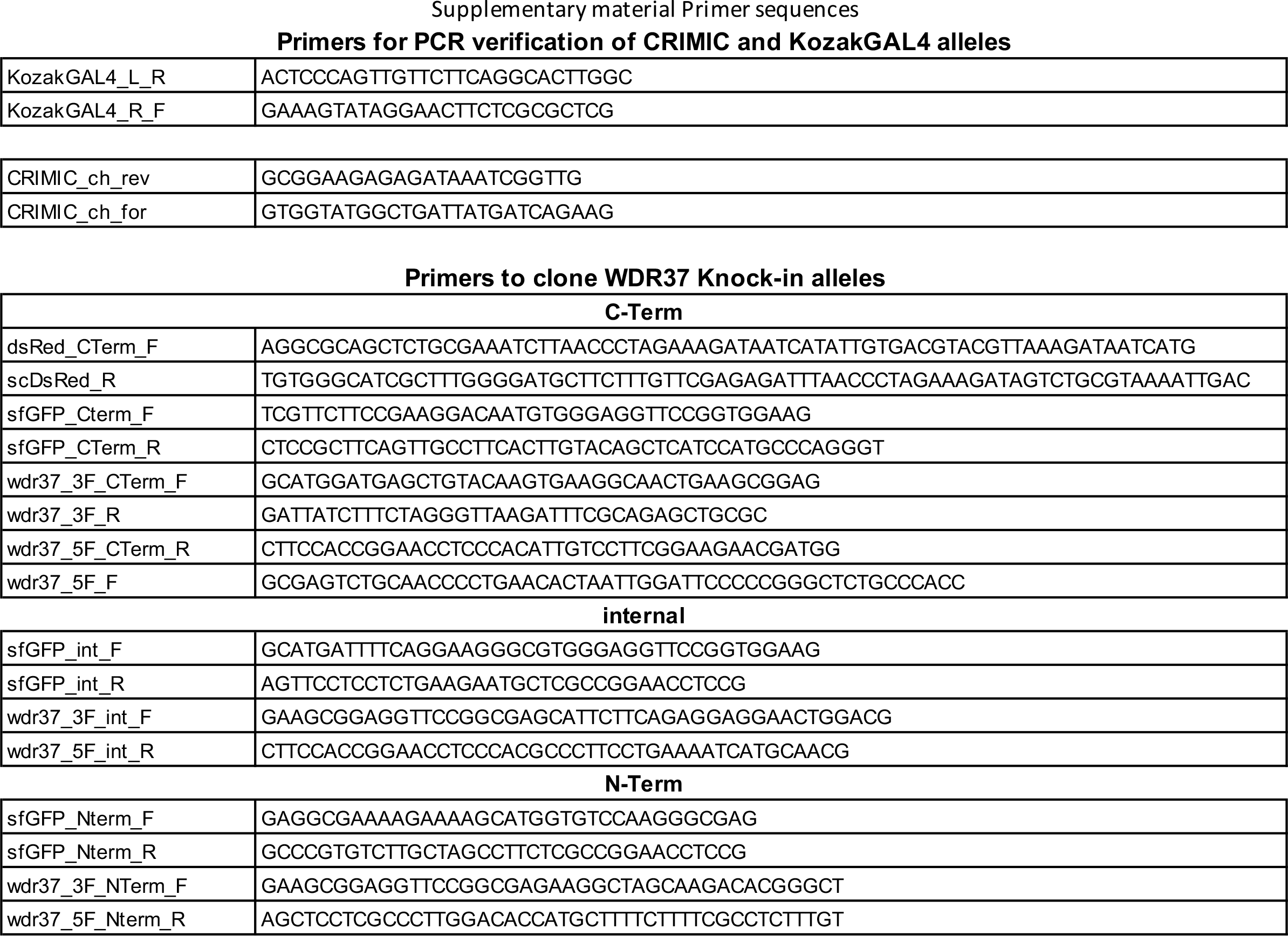
List of the 428 alleles generated in this study

## Supplementary methods Detailed cloning protocol and sequences

### Design of new CRIMIC synthesis constructs

Int200 strategy

Design region:

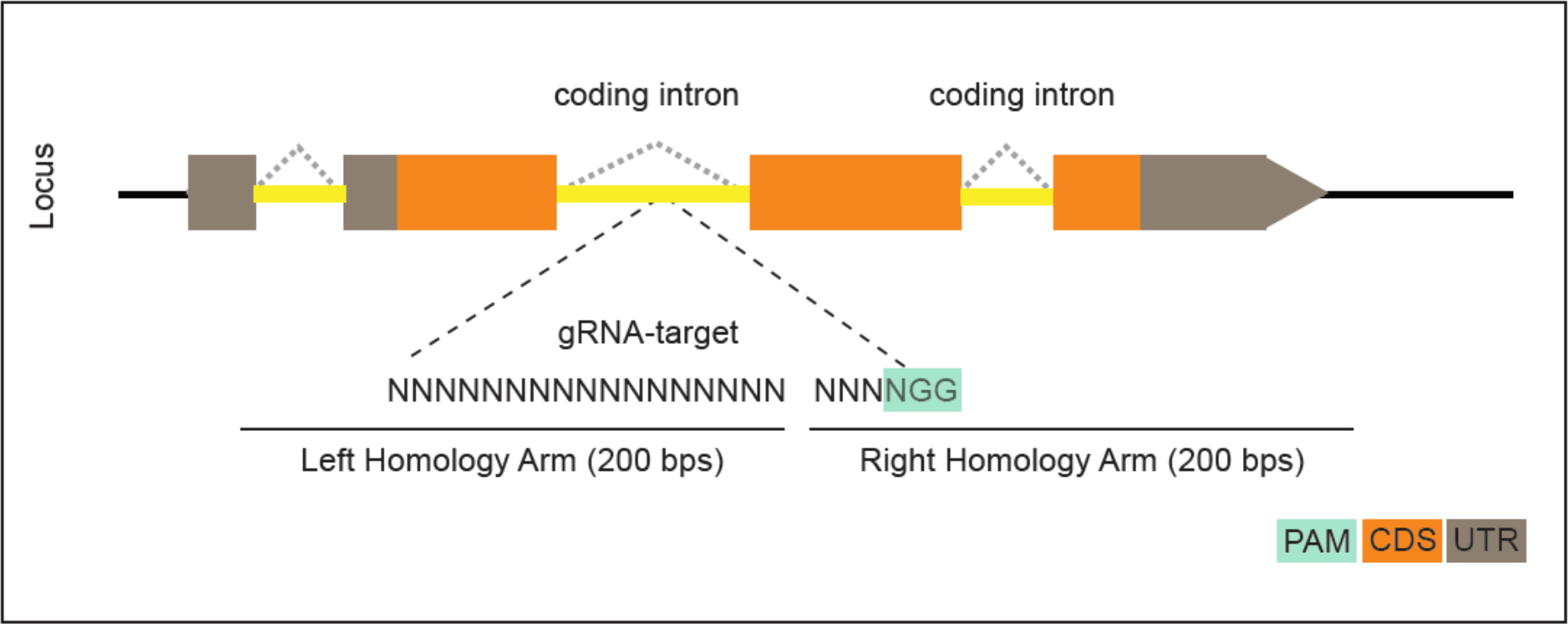

– 200 nucleotides on both sides of the cut site are used for synthesis using the template DNA file (int200_scaffold_BbsI file, in case the homology arms contain BbsI sites, BsaI-HF can be used for cloning. In that case int200_scaffold_BsaI scaffold file should be used for ordering the synthesis)

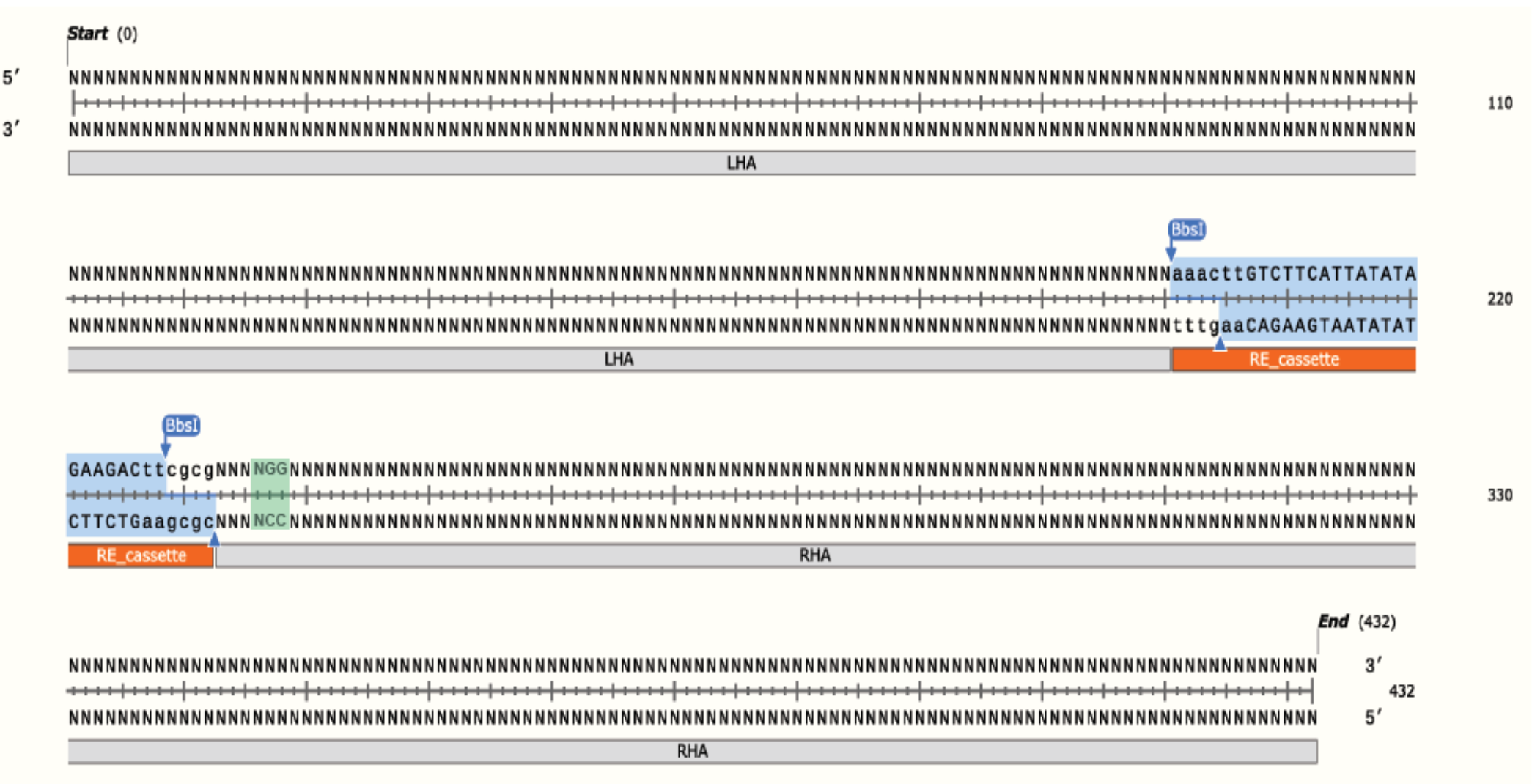

– Synthesis is ordered through Genewiz as Value gene synthesis (at 4nM scale). In order notes indicate:

- The constructs should be cloned in pUC57_Kan_gw_OK vector in EcoRV site. This vector has the gRNA1 target sites on either side of EcoRV cut site and hence places the homology arms in between gRNA1 cut sites. In the backbone there is a U6:gRNA1 that linearizes the construct *in vivo*.
- No need to order the glycerol stocks. Lyophilised constructs facilitates the reaction set up.

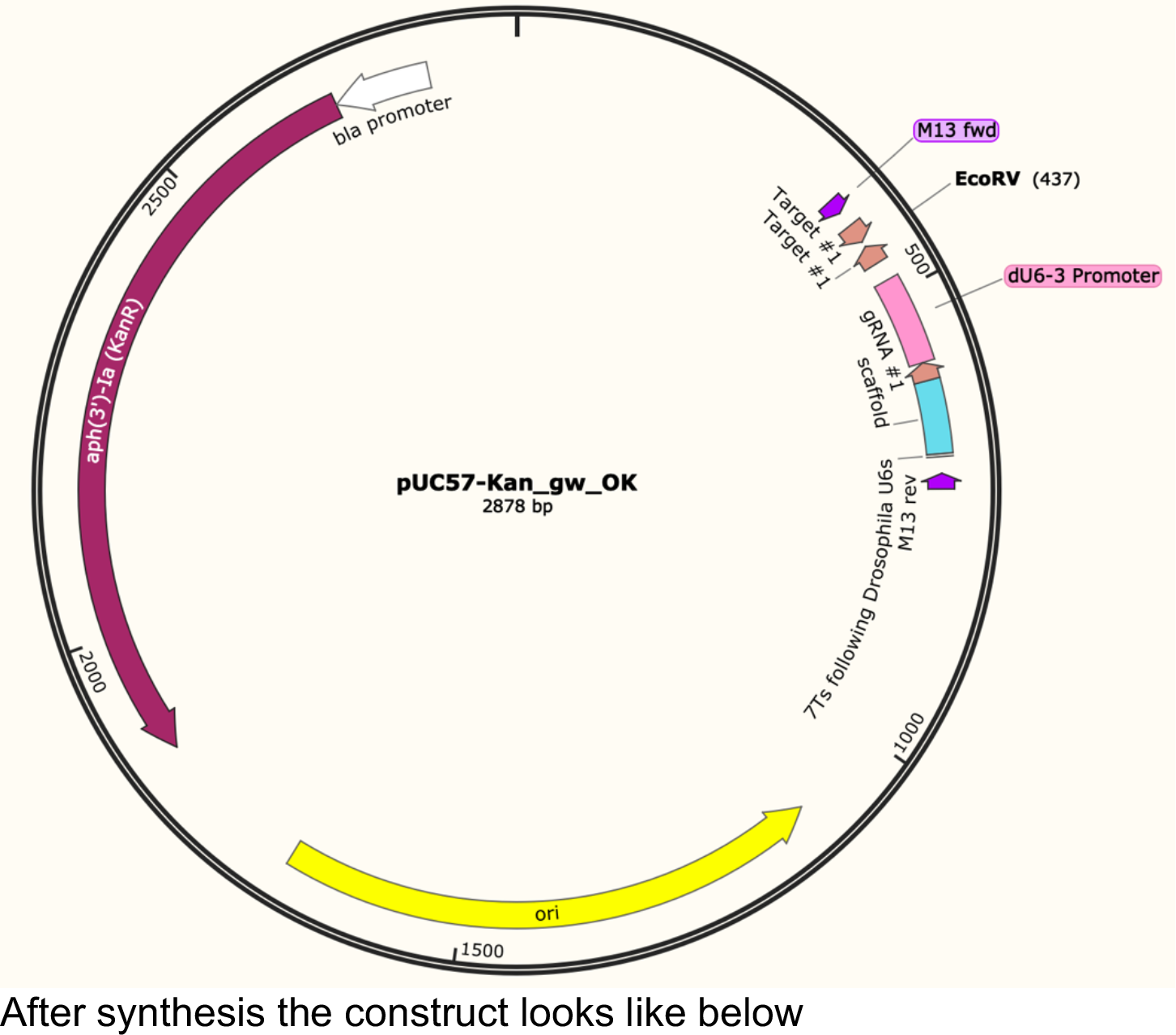

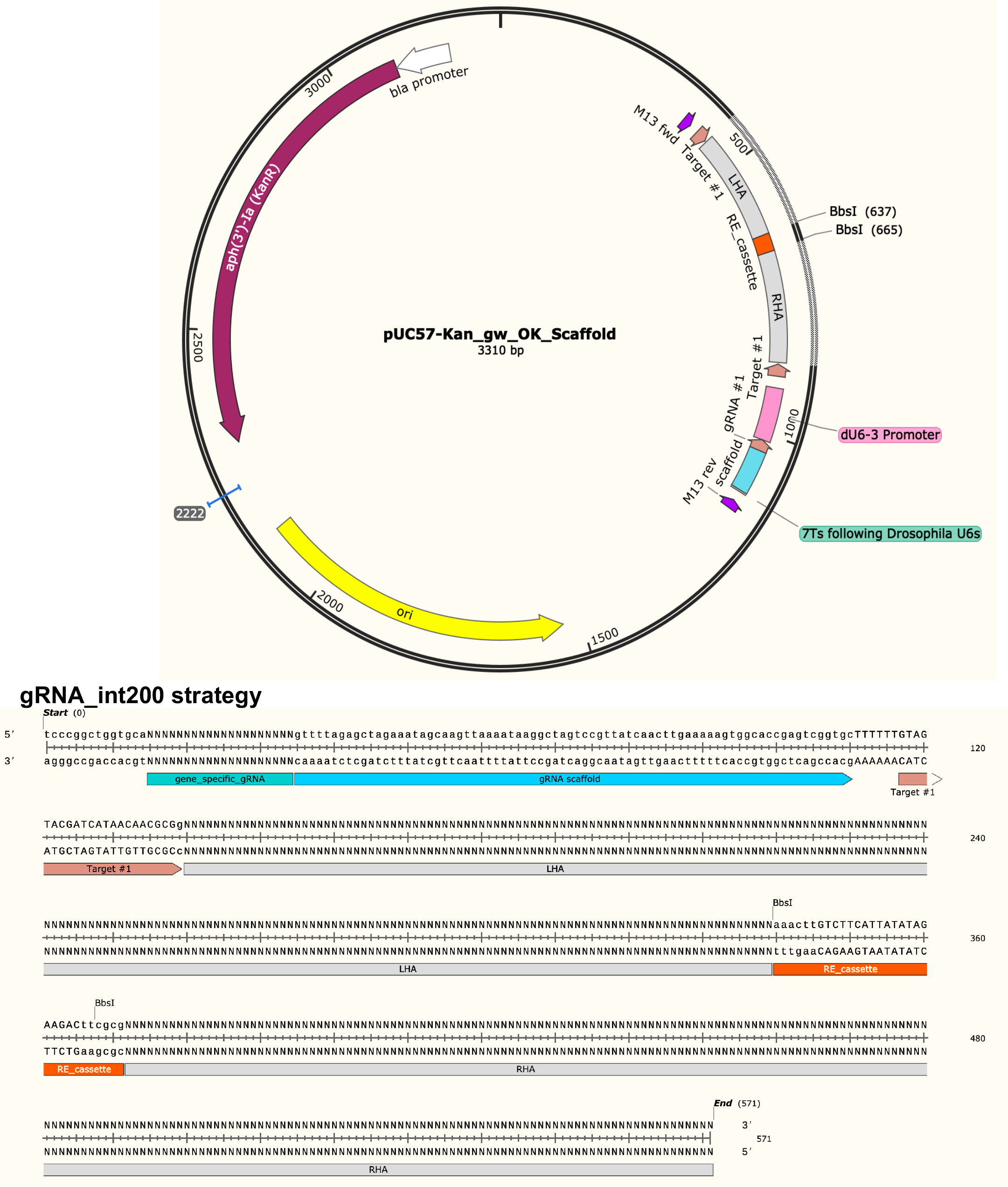

### gRNA_int200 strategy

The ordering for gRNA_int200 strategy is similar to int200 strategy with the exception of including the target specific sgRNA sequences in the synthesis order and the synthesis should be conducted on pUC57_Kan_gw_OK2 backbone that contains the rest of the components of the homology donor construct.

For *KozakGAL4* constructs 2 sgRNAs are added in the synthesis reaction.

### Cloning of constructs

– When the constructs arrive, resuspend them in 53µl dH2O (Do not resuspend in TE buffer. Genewiz lyophylises the constructs in TE buffer. Resuspending in TE buffer decreases efficacy of downstream cloning applications).
– Select the proper vector of pM37 (correct phase for T2AGAL4 or pM37_KozakGAL4 or pM37_SA_KozakGAL4) with BbsI-HF (or BsaI-HF if BsaI construct is being used). Set up the reaction (make a master mix for constructs of the same SIC if cloning multiple constructs):

- 1µl pM37-phase X* (290 ng/µl) or 1 µl pM37_KozakGAL4 (265 ng/µl)
- 2.5 µl 10X T4 DNA ligase buffer (NEB B0202S)
- 0.5 µl T4 DNA ligase (NEB M0202L)
- 1 µl Restriction enzyme (BbsI_HF (NEB R3733L) or BsaI_HFv2 (NEB R3559L)
- 19 µl of dH_2_O

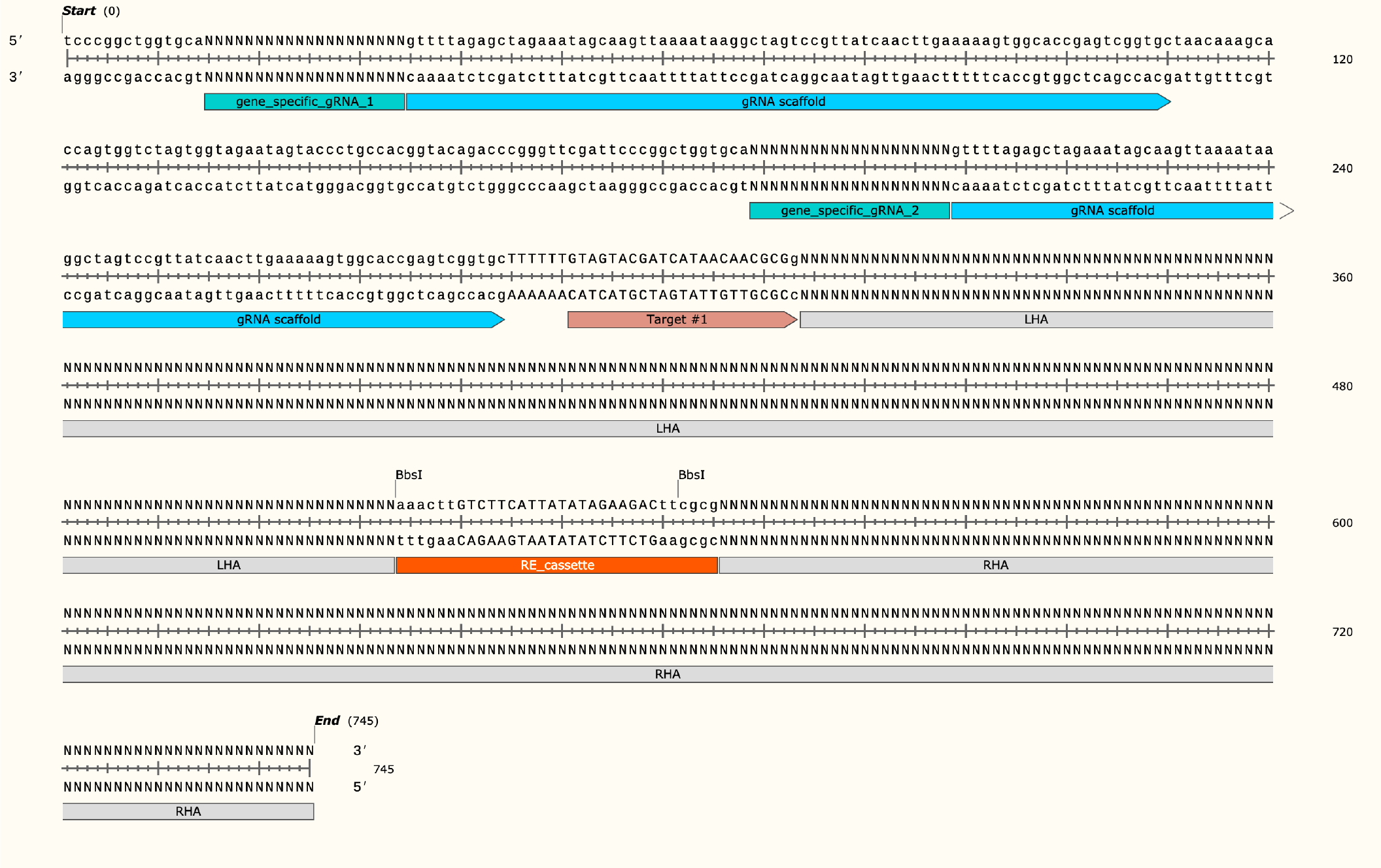

### Distribute the master mix in PCR tubes and add 1 µl of reconstituted intermediate plasmid

* Phase selection is done by following the reading frame of the gene at the end of the preceding exon. If the last codon in the preceeding exon is complete (NNN- intron-NNN), the phase is p0. If one of the nucleotide of the codon is in the preceeding exon and two are in the following exon (N-intron-NN) the phase is p1. If two nucleotides of the codon is in the preceeding exon and one in the following exon (NN-intron-N) the phase is p2. It is crucial to select the correct phase using incorrect phase cause a frameshift mutation.

-Incubate the reactions in a Thermocycler:

- 37°C 5 minutes
- 16°C 5 minutes
- Go to 1 30 times
- 65°C 20 minutes
- 8°C hold

The reactions can be left in the thermocycler overnight.

– An additional digestion step is done to remove self ligating plasmid backbones by adding:

- 19.5 µl dH2O
- 5 µl 10X Cutsmart buffer
- 0.5 µl BbsI or BsaI_HFv2 (the enzyme used for the cloning reaction)
- restriction mix can also be prepared as a mastermix and distributed to each sample 25µl/sample. Incukbate the reactions for 30 minutes in 37°C incubator or thermocycler.

– Transform to 50 ul chemocompetent DH5-alpha. Selection antibiotic is Kanamycin, hence 1-hour recovery is necessary after heatshock. Plate on LB plates with Kanamycin. Incubate at 37°C overnight.
– (Optional) Next day do colony PCR with primers M13F_Long_for CRIMIC_ch_rev M13F_long_for gacgttgtaaaacgacggccag
CRIMIC_ch_rev gcggaagagagataaatcggttg
I use an autoclaved micropipette tip to pick a colony, touch it on a gridded plate to copy the colony and dip the same pipette tip to PCR mix.

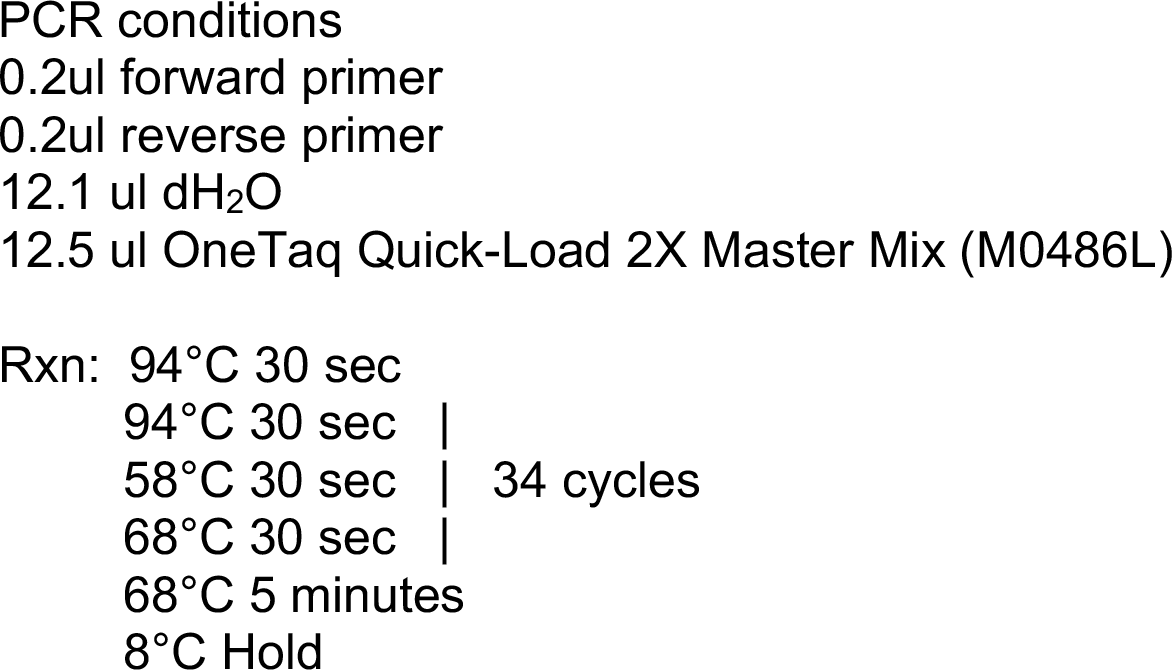

– The positive colonies will show ∼630 bps amplicon.
– The positive colonies are incubated in 5 ml of LB+Kanamycin and miiniprepped using Qiaprep Spin Miniprep kit (Qiagen 27106).
– Resulting DNA is sequenced using M13Reverse and intseq_forward primers (GTTCGATTCCCGGCCGATG)
Vector sequences:

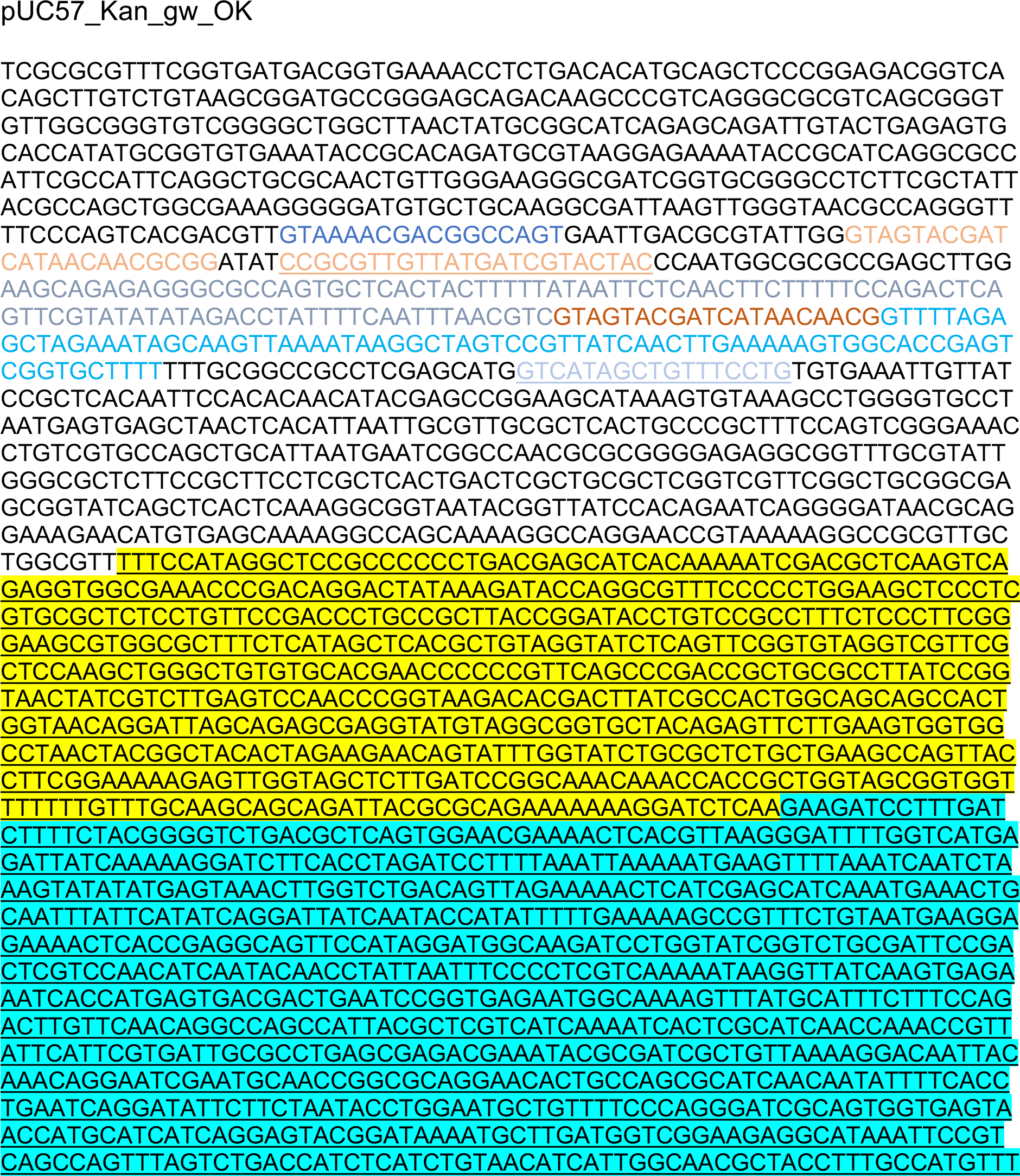
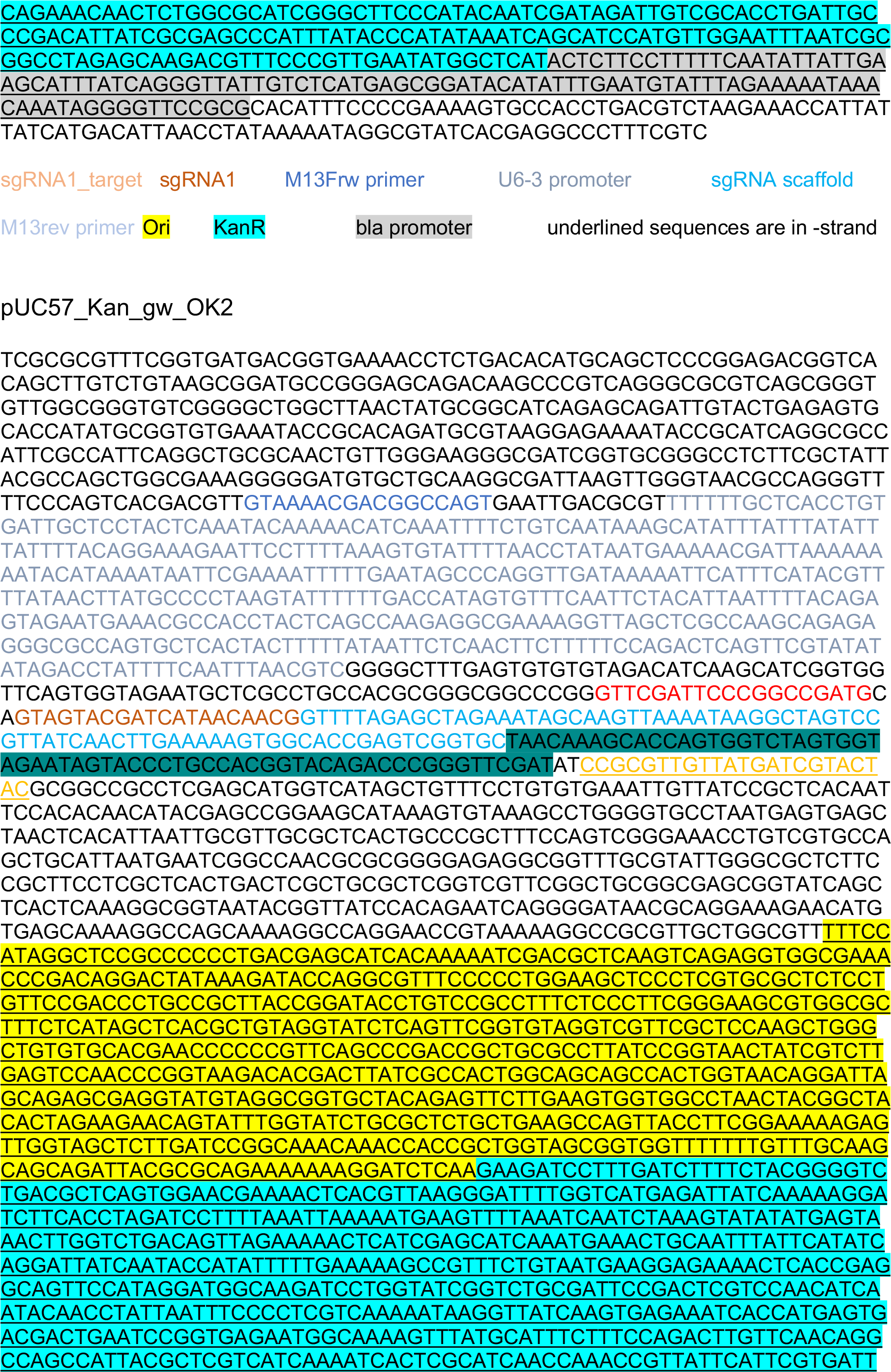
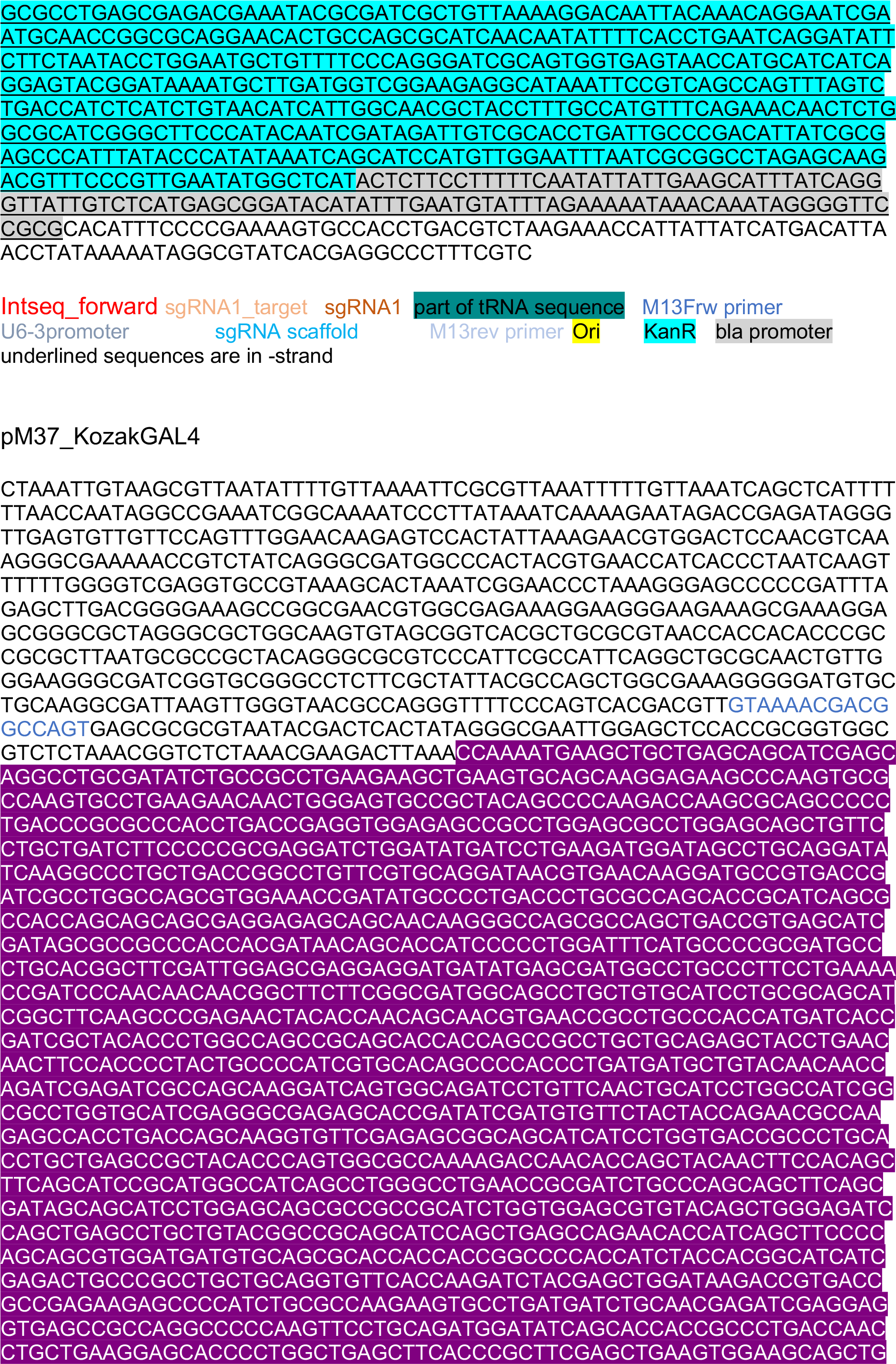
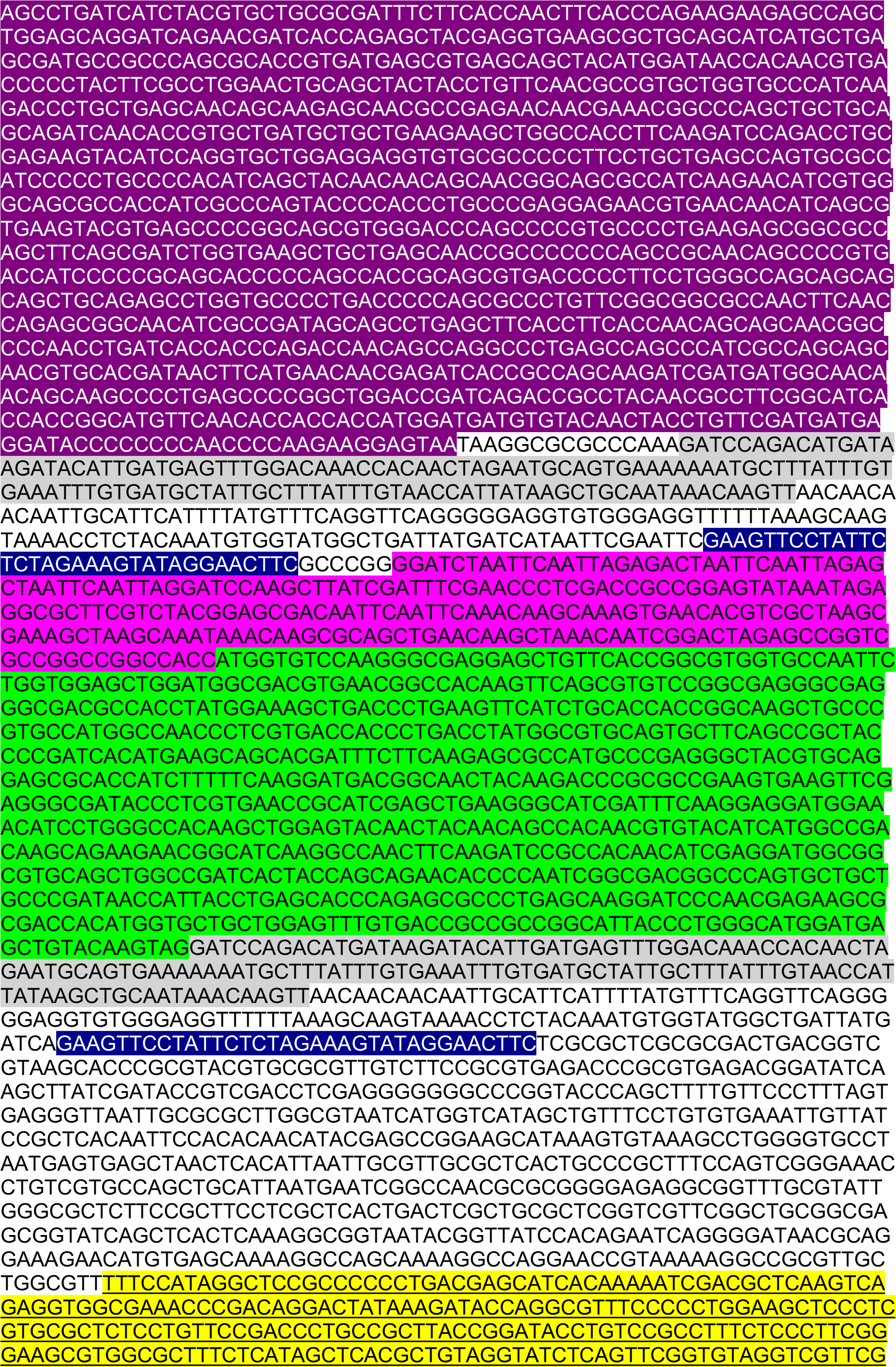
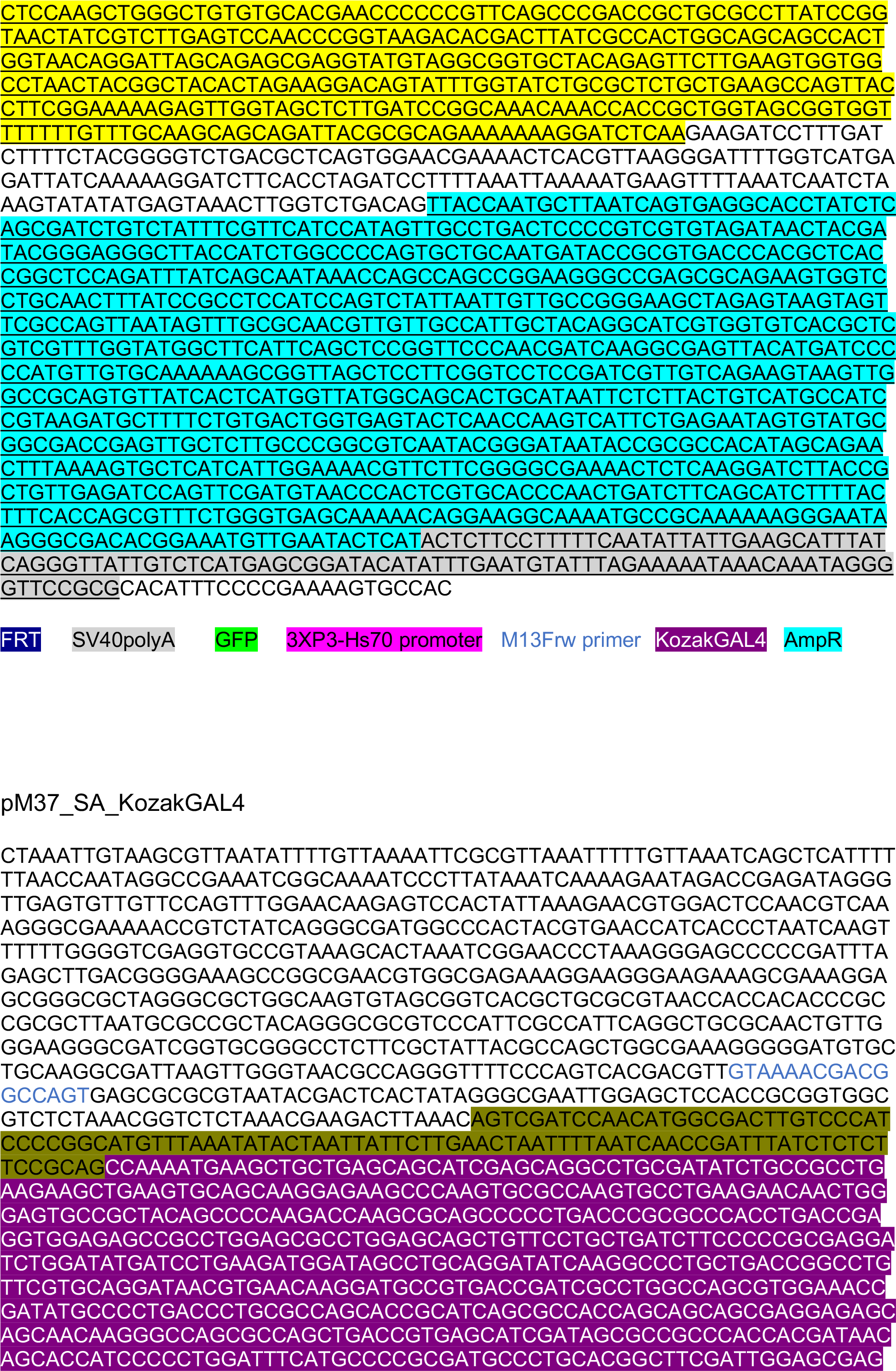
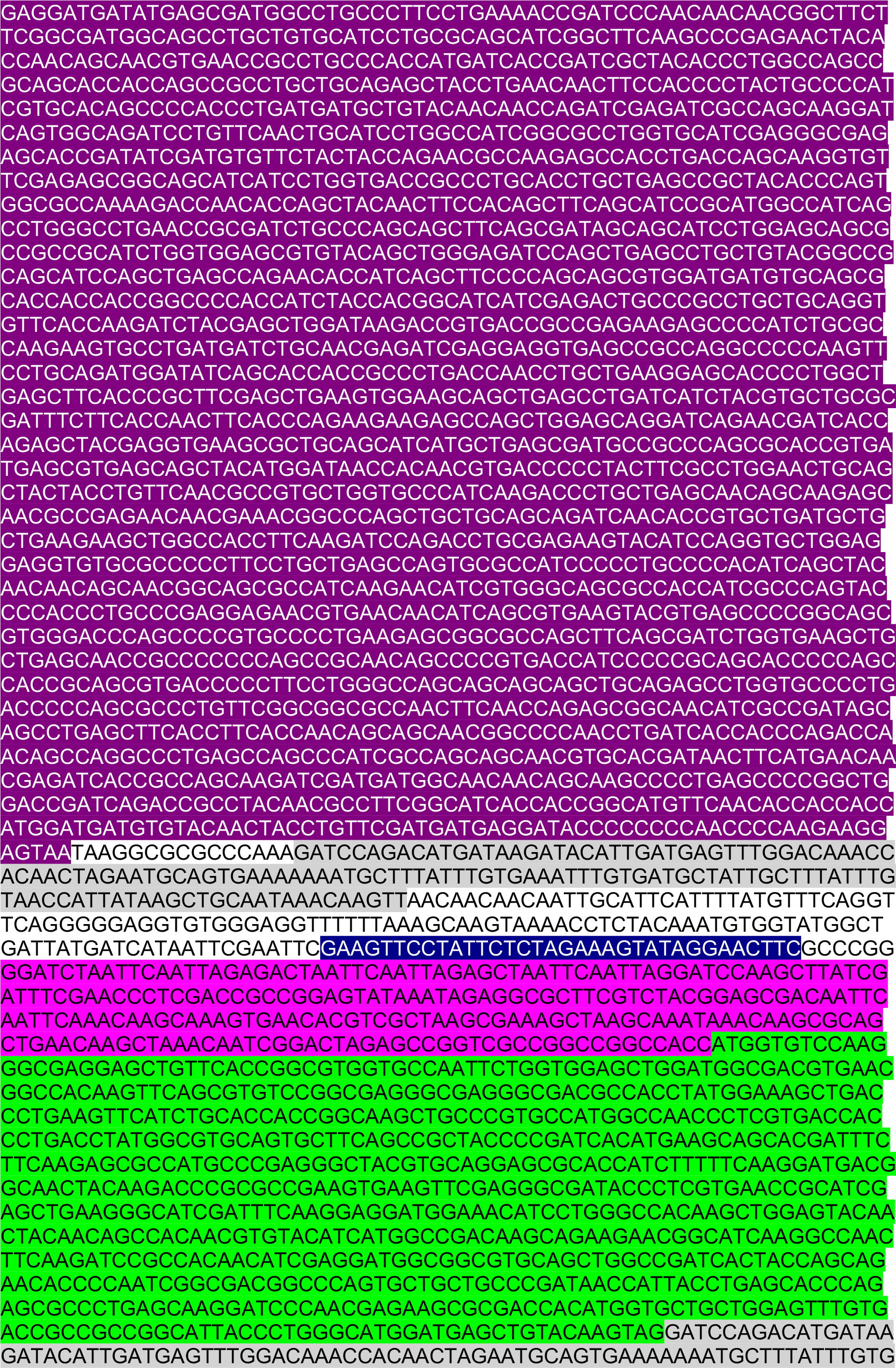
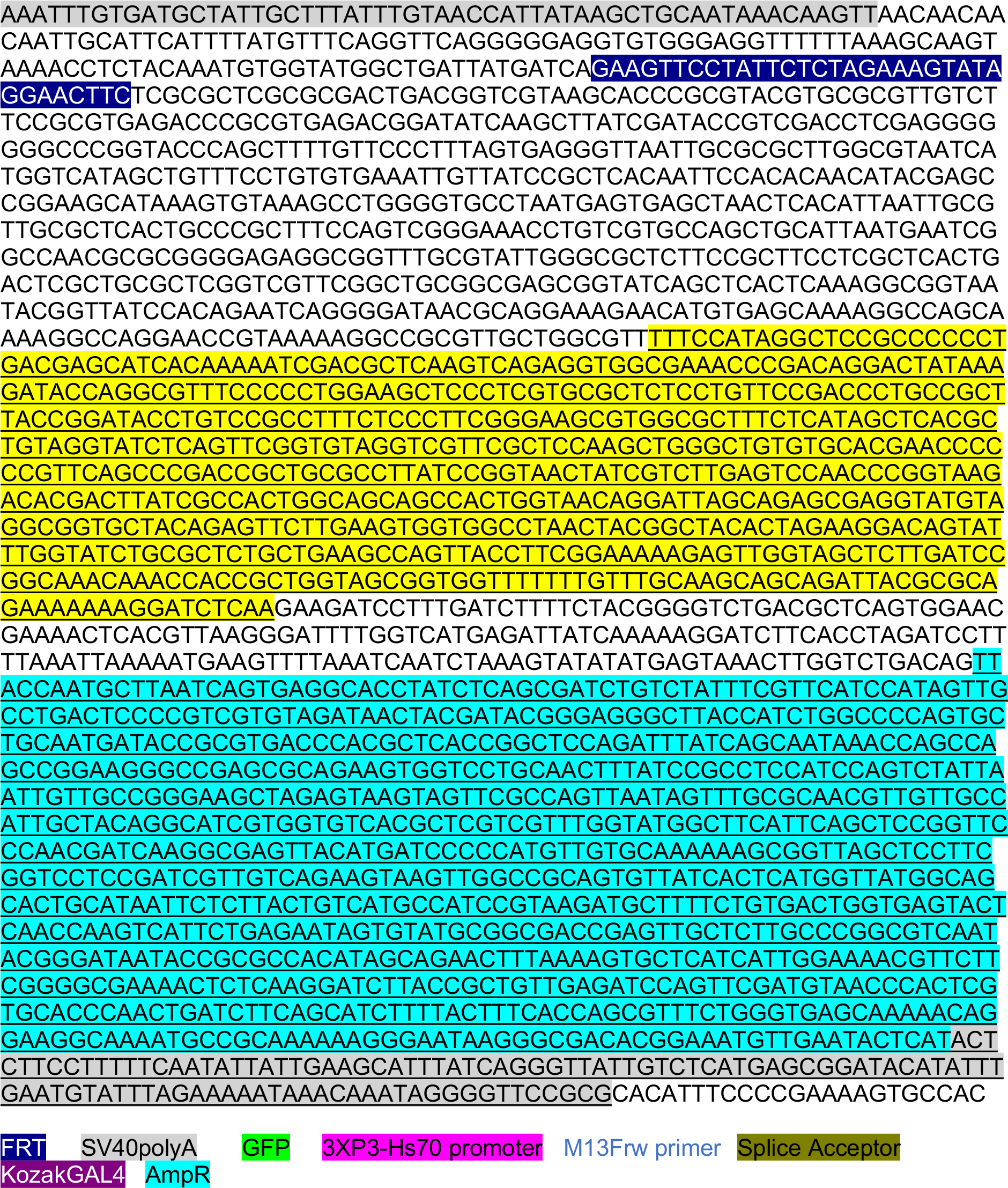

